# Neuregulin-1 regulates cardiomyocyte dynamics, cell cycle progression, and maturation during ventricular chamber morphogenesis

**DOI:** 10.1101/2022.11.28.518154

**Authors:** Joaquim Grego-Bessa, Paula Gómez-Apiñaniz, Belén Prados, Manuel José Gómez, Donal MacGrogan, José Luis de la Pompa

**Affiliations:** Centro Nacional de Investigaciones Cardiovasculares Carlos III (CNIC), Melchor Fernández Almagro 3, 28029 Madrid, SPAIN; CIBER de Enfermedades Cardiovasculares, Madrid, SPAIN; Bioinformatics Unit, CNIC, Madrid SPAIN

**Keywords:** Ventricular development, trabeculation, apico-basal polarity, oriented cell division, compaction, ventricular conduction system, actin cytoskeleton, EMT-like, cell cycle, Nrg1, ErbB2, Erk, Yap1

## Abstract

**BACKGROUND:** Cardiac ventricles are essential for providing the contractile force of the beating heart throughout life. How the primitive endocardium-layered myocardial projections called trabeculae form and mature into the adult ventricles is of great interest for fundamental biology and regenerative medicine. Trabeculation is dependent on the signaling protein Neuregulin-1 (Nrg1). However, the mechanism of action of Nrg1 and its role in ventricular wall maturation are poorly understood.

**METHODS:** In this study we investigated the functions and downstream mechanisms of Nrg1 signaling during ventricular chamber development using confocal imaging, transcriptomics, and biochemical approaches in mice with conditional cardiac-specific inactivation or overexpression of Nrg1.

**RESULTS:** Analysis of cardiac-specific-*Nrg1* mutant mice showed that the transcriptional program underlying cardiomyocyte-oriented cell division and trabeculae formation depends on endocardial Nrg1 to myocardial ErbB2 signaling and pErk activation. Early endothelial loss of Nrg1 and below normal pErk activation diminished cardiomyocyte Pard3 and Crumbs2 protein, and altered cytoskeletal gene expression and organization. These changes were associated with aberrant expression of genes involved in mitotic spindle organization and a directional shift from perpendicular to parallel/obliquely-oriented cardiomyocyte division. Further analysis indicated that Nrg1 is required for trabecular growth and ventricular wall thickening by regulating an epithelial-to-mesenchyme transition (EMT)-like process in cardiomyocytes involving migration, adhesion, cytoskeletal actin turnover, and timely progression through the cell cycle G2/M phase. Ectopic cardiac Nrg1 overexpression and high pErk signaling caused S-phase arrest, maintained high EMT-like gene expression and prolonged trabeculation, blocking compact myocardium maturation. Likewise, alterations of myocardial trabecular patterning resulting from above– or below-normal Nrg1-dependent pErk activation were concomitant with disorganization of the sarcomere actin cytoskeleton. The Nrg1 loss– and gain-of-function transcriptomes were enriched for yes-associated protein-1 (Yap1) gene signatures, identifying Yap1 as a potential downstream effector. Biochemical and imaging data showed that pErk activation and nuclear-cytoplasmic distribution of Yap1 during trabeculation are dependent on Nrg1.

**CONCLUSIONS:** These data establish the Nrg1-ErbB2/4-pErk axis as a crucial regulator of cardiomyocyte cell cycle progression and migration during ventricular development. Moreover, our data identify a Nrg1-dependent signaling cascade that could be leveraged for future cardiac regenerative therapies.

**Novelty and Significance:** *WHAT IS KNOWN?:*

- Myocardial trabeculae play important roles in ventricular chamber growth, development of the conduction system, and formation of the coronary arteries.
- Trabeculae are formed through oriented cell division (OCD), and their growth is driven by directional migration.
- The membrane glycoprotein Neuregulin-1 (Nrg1) mediates cell-cell signaling and is essential for trabecular development.

*WHAT NEW INFORMATION DOES THIS ARTICLE CONTRIBUTE?:*

- Nrg1 signaling is essential for the expression of cardiomyocyte polarity genes and the organization of the cytoskeleton during the oriented cell division process underlying trabeculation.
- Nrg1 is required for the formation of the inner ventricular wall but not the coronaries.
- Nrg1 regulates motility and cell-cycle progression during ventricular wall growth.
- Ectopic expression of Nrg1 leads to excessive trabeculation of the myocardium and disrupts compaction.
- Nrg1 regulates ventricular patterning mediated by cytoskeletal dynamics and modulates pErk-dependent Yap1 S274 phosphorylation during trabeculation.
- Nrg1 is not required for ventricular compaction.

## Introduction

Ventricular chamber development entails transient morphogenetic changes that include the formation of trabeculae, their expansion and growth, and their subsequent incorporation into a compact muscular wall. Fate-mapping and time-lapse live imaging in zebrafish has shown that trabeculae form by the extrusion of cardiomyocytes and their radial extension along the inner curvature of the looping heart and into the ventricular lumen ^1–5^. In mice, trabeculation begins with oriented cell division (OCD) perpendicular to the ventricular inner wall plane, and trabecular growth is driven by directional migration from the outer (epicardial) layer toward the inner (endocardial) layers ^6–8^. The process of trabeculae coalescence forms a dense meshwork that maximizes ventricular surface area for nutrient and gas exchange in the absence of a coronary circulation ^9^. This meshwork later differentiates into a specialized ventricular conduction system and is resolved by expansion of the compact myocardial wall, with trapped endocardial cells giving rise to the coronary arteries ^10–14^.

Trabeculation defects or failed compaction in humans, manifests as hypertrabeculation or non-compaction phenotypes ^15^. Trabeculae thickening, decreased intertrabecular spacing, and underdevelopment of the compact wall myocardium are defining morphological features of left ventricular non-compaction cardiomyopathy ^16^. This phenotype is observed in various conditions, that have been associated with sarcomeric genes, cardiac contractility, calcium handling, and developmental signaling pathways ^17^. Conversely, acquired non-compaction or the appearance of hypertrabeculation in trained athletes or during pregnancy, have been interpreted as morphophysiological manifestations of increased hemodynamic load ^18^. Moreover, genetic association and Mendelian randomization studies have found an inverse relationship between the complexity of trabecular morphology and the risk of cardiovascular disease, pointing to a potentially beneficial role of excessive trabeculation in adults ^19^. Given the diversity of underlying pathophysiological mechanisms, there is ongoing debate about the significance of hypertrabeculation and non-compaction within the spectrum of myocardial adaptive processes and diseases ^18, 20–22^.

Trabecular development requires the intercellular signaling action of the membrane glycoprotein Neuregulin-1 (Nrg1) ^4, 23–25^. This endocardial protein directly binds to epidermal growth factor receptor tyrosine kinase (ErbB4) on adjacent cardiomyocytes, triggering ErbB2 ligand-stimulated heterodimerization and tyrosine phosphorylation, followed by activation of downstream pathways ^26^, including the phospho-extracellular signal-regulated kinase (pErk) intracellular signaling pathway ^27^. Targeted mutation of *Erbb2* or *nrg2a* in fish ^3, 4, 25^, or deletion of *Nrg1*, *ErbB2*, or *ErbB4* in mice ^24, 28–32^, blocks trabeculation. Cardiomyocyte proliferation and contractility, which are required for trabeculation, are impaired in mutant *ErbB4* mice ^33^ and *erbb2* zebrafish ^3^, whereas potentiation of ErbB2 signaling in mouse adult cardiomyocytes promotes proliferation and myocardial regeneration ^34^. Consistent with these findings, beneficial effects of Nrg1 for cardiac function have been reported in mouse models of cardiac injury ^33^; explanted human tissue ^35^; ischemic, dilated, and viral cardiomyopathy ^36^; and heart failure ^37–39^. Clinical trials with different forms of recombinant Nrg1 support a cardioprotective role in systolic heart failure ^40^, although the target cell-type-specific effects remain to be characterized.

The essential role of Nrg1 in the initiation of trabeculation is well documented, but its role in the cellular and molecular basis of trabecular growth, maturation, and resolution remains uncertain. In particular, understanding is limited about how morphological transitions occur and progress to yield a thick ventricular wall. Mutations affecting trabeculation in mice invariably lead to fetal demise ^41^, precluding direct analysis of the mechanism(s) of trabecular growth and maturation.

Here, we used conditional and temporally regulated cardiac loss– and gain– of function mutants and gene profiling to show that pErk activation by Nrg1 promotes trabeculation by sustaining a transcriptional program regulating cell polarity, mitotic spindle orientation and cytoskeletal dynamics thereby facilitating trabecular growth and maturation. Additionally, Nrg1-induced pErk activation drives dynamic changes in cardiomyocyte shape and movement, leading to trabecular expansion and thickening of the ventricular wall. Furthermore, this signaling cascade plays a role in timely progression through the cell cycle. Notably, we identified significant overlap between Nrg1-dependent genes and conserved gene signatures regulated by the ErbB2 receptor and the core component of the MuvB-complex, Lin9. Both of these factors act upstream or in conjunction with the Hippo pathway, which involves yes-associated protein-1 (Yap1). These findings suggest a complex interplay between Nrg1-ErbB2/4-Erk and Hippo-Yap1 signaling, which holds promise for potential cardiac regeneration therapies in the future.

## METHODS

Full *Methods* are provided in the *Online Data Supplement*. Animal studies were approved by the CNIC Animal Experimentation Ethics Committee and by the Community of Madrid (Ref. PROEX 155.7/20). All animal procedures conformed to EU Directive 2010/63EU and Recommendation 2007/526/EC regarding the protection of animals used for experimental and other scientific purposes, enacted in Spanish law under Real Decreto 1201/2005.

## RESULTS

### Nrg1-ErbB2/4-pErk signaling is required for trabeculae formation and expansion

To study the role of Nrg1-ErbB2/4 signaling in cardiac development, we used a conditional *Nrg1^flox^* mouse line ^42^ (see *Methods*). We crossed *Nrg1^flox^*mice with the *Tie2^Cre^* driver line ^43^ to conditionally inactivate *Nrg1* signaling in endothelial and endocardial cells from E7.5 onwards. At E8.5, there were no overt differences between control and *Nrg1^flox^*;*Tie2^Cre^* embryos, both featuring a bilayered myocardial epithelium (Figure S1A,C). At E9.5, control hearts already showed nascent trabeculae and a double-layered compact wall (Figure 1A; Figure S1B). In contrast, *Nrg1^flox^*;*Tie2^Cre^* mutants contained only a few trabecular projections, and the ventricular wall appeared somewhat thicker (Figure 1B,C; Figure S1D). By E10.5, the trabecular network in control ventricles had become more complex, whereas *Nrg1^flox^*;*Tie2^Cre^* mutants had only a few thick, rudimentary trabeculae and a noticeably thicker compact myocardium, especially in the left ventricle (Figure S1E,F,G,H). *Nrg1^flox^*;*Tie2^Cre^* mutants die by E11.5-12.5 (Suppl. Table S1, sheet 1). The profound differences in compact wall thickness and trabecular network size and complexity were also revealed by 3D-reconstructions (Figure 1D,E,F; Figure S1I,J; Movies S1 and S2). This defective trabecular phenotype was broadly reproduced in *Nrg1^flox^* mice crossed with the cardiac mesoderm driver line *Mesp1^Cre^* ^44^ (Figure S2A-D; Suppl. Table S1, sheet 2).

**Figure 1.**
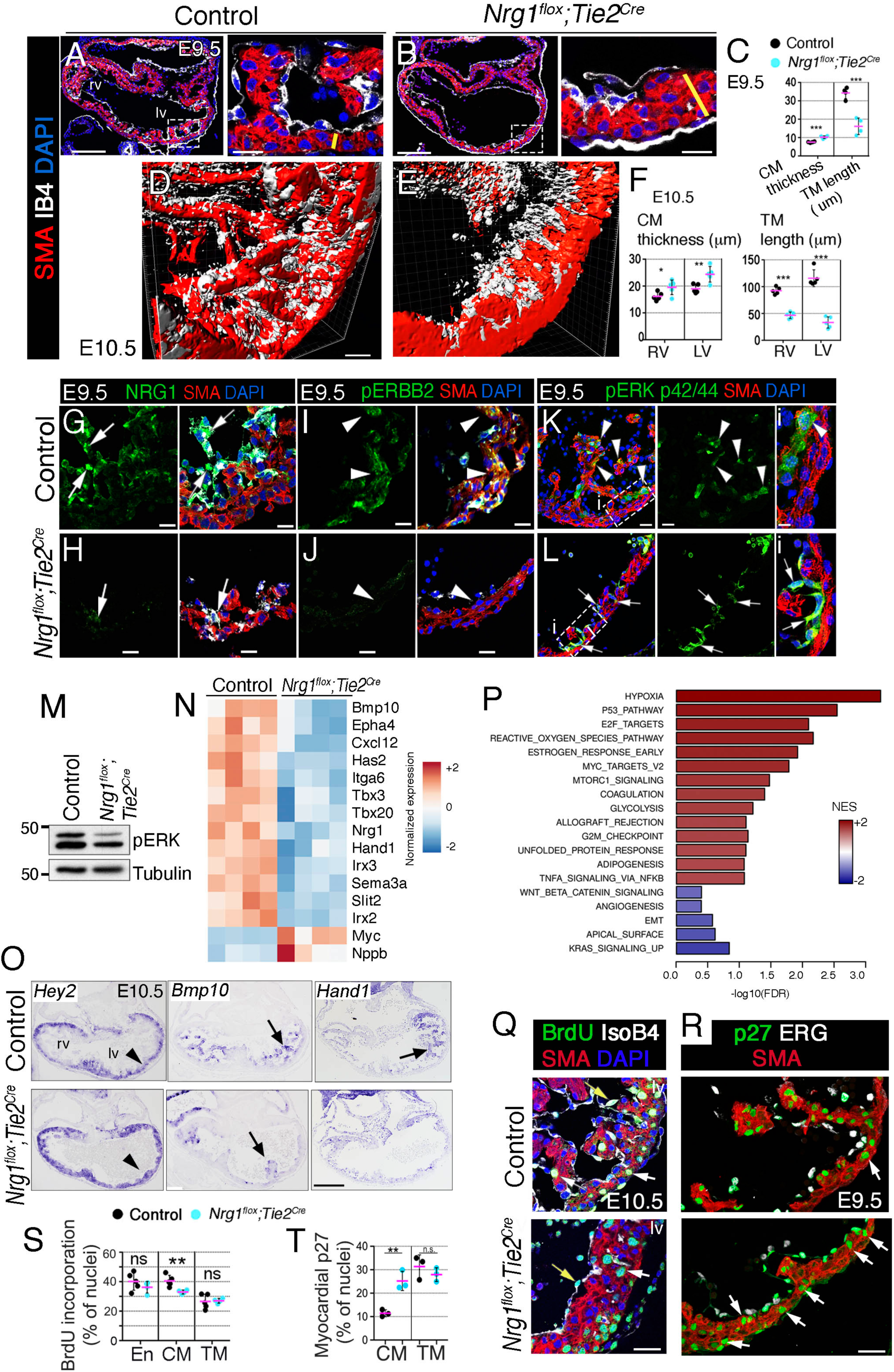
Nrg1-ErbB2-pErk signaling is required for trabeculation initiation and expansion. (A,B) Immunofluorescence against smooth muscle actin (SMA, red) and isolectin B4 (IB4, white) in heart sections from E9.5 control **(A)** and *Nrg1^flox^;Tie2^Cre^* embryos **(B)**. Yellow lines mark the thickness of the compact myocardium (CM). Sections were counterstained with DAPI (blue). **(C)** Quantification of CM thickness and trabecular myocardium (TM) length in control and *Nrg1^flox^;Tie2^Cre^* left ventricles at E9.5. **(D, E)** 3D reconstruction of SMA-stained E10.5 control and *Nrg1^flox^;Tie2^Cre^* left ventricles. **(F)** Quantifications of CM thickness and TM length in control and *Nrg1^flox^;Tie2^Cre^* ventricles at E10.5. **(G-L)** Immunofluorescence against Nrg1, pErbB2, and pErk (green) in E9.5 control embryos **(G, I, K)** and *Nrg1^flox^;Tie2^Cre^* embryos **(H, J, L)**. SMA, red; IB4, white; DAPI counterstain, blue. Arrows mark endocardium, arrowheads mark myocardium. **(M)** Representative western blot of pErk in E9.5 ventricles (pool of n = 3 per genotype). **(N)** Heatmap of representative differentially expressed genes (DEGs) in E9.5 control vs. *Nrg1^flox^;Tie2^Cre^*hearts. The color code represents normalized gene expression from –2 to +2. **(O)** ISH of *Hey2*, *Bmp10*, and *Hand1* in E9.5 control and *Nrg1^flox^;Tie2^Cre^* embryos. Arrows mark TM, arrowheads CM. **(P)** Hallmark Gene Set Enrichment Analysis (GSEA) for E9.5 *Nrg1^flox^;Tie2^Cre^* vs. Control. The bar plot represents enrichment data for 19 Hallmark gene sets at a false discovery rate FDR qval < 0.1 and FDR qval < 0.5, for gene sets with positive or negative enrichment score, respectively. Of those gene sets, 8 had an FDR qval < 0.05. The scale bar represents normalized enrichment score (NES) from –2 to +2. Positive scores (red), indicating enrichment in *Nrg1^flox^;Tie2^Cre^*relative to the control condition, were found for 14 gene sets, and negative scores (blue), indicating depletion, were found for 5 gene sets. Relative expression values for the “leading edge” genes associated to gene sets P53_PATHWAY, E2F_TARGETS, KRAS_SIGNALING_UP and EMT are presented as heatmaps in Figure S4. **(Q)** BrdU immunodetection in E10.5 control and *Nrg1^flox^;Tie2^Cre^*embryos. Arrows mark myocardium, arrowheads endocardium. **(R)** p27 immunodetection in E9.5 control and *Nrg1^flox^;Tie2^Cre^* embryos. Arrows mark myocardium. **(S)** Quantification of BrdU incorporation in E10.5 control embryos (n=4) and *Nrg1^flox^;Tie2^Cre^* embryos (n=3). **(T)** Quantification of p27 immunostaining in E9.5 control and *Nrg1^flox^;Tie2^Cre^*embryos (n=3 per genotype). En, endocardium; RV, right ventricle; LV, left ventricle. * *P* < 0.05, ** *P* < 0.01, *P* < 0.001; ns, non-significant by Student t-test. Scale bars, 200 μm (low magnification in A,B, and O); 50μm (D,E); 20 μm (high magnification in A,B and G, I, K H, J, L, Q, and R); 10 μm (high magnification in Gi,Hi).

E9.5 *Nrg1^flox^*;*Tie2^Cre^* mutant hearts showed severely attenuated endocardial Nrg1 expression (Figure 1G,H). This was reflected in reduced Nrg1 signaling activity in the trabecular myocardium at E9.5, indicated by weak myocardial expression of phosphorylated ErbB2 (pErbB2) (Figure 1I,J) and of the phosphorylated intracellular effector Erk (pErk) (Figure 1K,L,M).

We examined *Nrg1* transcription at key developmental stages by *in situ* hybridization (ISH) (Figure S2E-I). At E10.5 *Nrg1* was transcribed robustly and uniformly in the endocardium (Figure S2E), and by E12.5 and until E16.5 at least, Nrg1 transcripts persisted in scattered cells throughout the endocardium (Figure S2F-I). We further assessed pErk expression at key cardiac developmental stages as a readout of Nrg1-pErbB2 activity ^27, 34, 45^ (Figure S2J-N). At E9.5, intense but patchy pErk signals were detected in the myocardium, especially at the tips of trabeculae, but not in endocardium (Figure S2J). At E11.5, pErk expression was maintained along the trabeculae, especially at the tips (Figure S2K). At E12.5, strong pErk expression was found at the base of trabeculae (Figure S2L), suggesting a role in the formation of endocardium-derived coronary vessels. At E13.5, weak pErk expression was detected in a few myocardial cells, and stronger expression in the nascent coronaries (Figure S2M). At E16.5, pErk expression was almost exclusively limited to the coronary endothelium (Figure S2N). The pErk temporal expression data are consistent with a function of Nrg1 in trabecular myocardium between E9.5 and E12.5, and in coronary endothelium from E13.5 onwards.

### Nrg1 is required for cardiac development patterning and proliferation

To understand the basis of the defective trabeculation in *Nrg1^flox^*;*Tie2^Cre^*mutants, we examined the extracellular matrix (ECM) between endocardium and myocardium (the cardiac jelly), which is required for directional migration and the formation of cardiac trabeculae ^28, 46^. In control and *Nrg1^flox^*;*Tie2^Cre^* hearts at E8.5, Alcian blue staining revealed the glycosaminoglycan constituent of the cardiac jelly (Figure S3A,C). However, at E9.5 and more obviously at E10.5, Alcian blue-staining was severely diminished in *Nrg1^flox^*;*Tie2^Cre^* hearts compared with controls (Figure S3B,D,E,F), consistent with the manifestation of the trabeculation defect.

We next conducted an RNA sequencing (RNA-seq) analysis of E9.5 control and *Nrg1^flox^*;*Tie2^Cre^* hearts. We found 1,219 differentially expressed genes (DEGs; 651 upregulated and 515 downregulated; *P<*0.05; Table S2, sheets 1 and 2). *Nrg1^flox^*;*Tie2^Cre^* heart sections showed severely depleted expression of *Nrg1* and several key cardiac developmental genes, including the trabecular marker genes *Bmp10, Hand1, Irx3,* and *Sema3a* (Figure 1N; Figure S3G). Also decreased in *Nrg1^flox^*;*Tie2^Cre^* mutants was the expression of *Has2*, which participates in the synthesis of the cardiac jelly ^46^ (Figure 1N). ISH showed that *Has2* was restricted to a few cells in the rudimentary trabeculae of *Nrg1^flox^*;*Tie2^Cre^* mutants (Figure S3G,H), as was the expression of *Bmp10*, *Hand1, Irx3,* and *Sema3a* (Figure 1O; Figure S3I-L). In further agreement, *Nrg1^flox^*;*Tie2^Cre^*mutants had markedly depleted expression of integrin alpha 6 (ITGα6) (Figure S3M-O), a trabeculae-restricted integrin ^47^. In contrast, ISH revealed increased *Hey2* expression in the thicker compact myocardium of *Nrg1^flox^*;*Tie2^Cre^*mutants (Figure 1O). These results indicate that abrogation of Nrg1-ErbB2/4 signaling disrupts trabeculae formation and chamber myocardium patterning.

To define mechanisms, we performed a GSEA of “HALLMARK” gene sets ^48^. Most pathways were enriched (FDR q-val <0.05), including cellular stress pathways (HYPOXIA and REACTIVE_OXYGEN_SPECIES_PATHWAY), associated with activation of the p53 (P53_PATHWAY; Figure S4A). The p53 protein responds to stresses that disrupt the fidelity of DNA replication and cell division and initiates a program of cell cycle arrest, cellular senescence, or apoptosis ^49^. Hence, the genes contributing most to the enrichment score (the leading-edge genes) of the P53_PATHWAY were associated with the cell cycle, DNA damage response, apoptosis, and other processes typical of the p53 response (Figure S4A). Other enriched pathways were associated with proliferation and growth (ESTROGEN_RESPONSE_EARLY, MYC_TARGETS_V2, MTORC1_SIGNALING, E2F_TARGETS) (Figure 1P; Table S2, sheet 3). E2F_TARGETS represent genes involved in the cell cycle, chromatin homeostasis, DNA repair, DNA replication, mitosis, and other processes, including cytoskeletal regulation (Figure S4B).

To define the nature of the proliferative defect in *Nrg1^flox^*;*Tie2^Cre^*hearts, we measured the incorporation of bromodeoxyuridine (BrdU, an S-phase marker) from E8.5 and E10.5 (Figure 1Q; Figure S5). At E8.5, the proliferation rate in mutant hearts was similar to that of control hearts (Figure S5A,B). At E9.5, a statistically significant 10% relative reduction in proliferation was detected in the compact myocardium of *Nrg1^flox^*;*Tie2^Cre^* hearts (50% vs. 40%; Figure S5C-E). At E10.5, this difference persisted in compact myocardium (Figure 1Q,S), whereas proliferation in the endocardium and trabecular myocardium was unchanged (Figure 1Q,S). Closer inspection of the E2F_TARGET “leading edge” genes further revealed that many cell cycle inhibitors including p21/*Cdkn1a,* p27/ *Cdkn1b,* p18*/Cdkn2c,* CIP2*/Cdkn3,* and were upregulated (Figure S4). *Cdkn1b* encodes for cyclin-dependent kinase inhibitor p27^Kip1^ protein which binds and blocks cyclin-CDK to arrest the cell cycle ^50^ and organismal growth ^51, 52^. Consistent with reduced BrdU measurements, p27 protein expression was readily increased in *Nrg1^flox^*;*Tie2^Cre^*compact myocardium, but did not change in trabecular myocardium (Figure 1R,T).

*Nrg1^flox^*;*Tie2^Cre^* hearts were also characterized by altered pathways previously linked to trabeculation in zebrafish ^1, 5, 53^; namely, KRAS_SIGNALING_UP, APICAL_SURFACE, EMT (EPITHELIAL_MESENCHYMAL_TRANSITION) (Figure 1P; Figure S4C,D; Table S2, sheet 3). These processes, involving the cytoskeleton, cellular motility and the ECM tended to be depleted although this change was only evoked (FDR qval<0.5). (Figure1P, Figure S4C,D). We surmise that cardiomyocytes that typically contribute to trabeculae formation through migratory processes, remain in the compact layer of *Nrg1^flox^*;*Tie2^Cre^*mutants, leading to abnormal ventricular thickening at E9.5-10.5, despite the attenuated proliferation. Overall, our data indicate that Nrg1-ErbB2/4 signaling is required for both the patterning and emergence of trabeculae, and the proliferation of compact layer cardiomyocytes.

### Nrg1 is required for polarity gene expression and cytoskeletal organization during trabeculation

We further inspected the RNA-seq data for changes in genes associated with cell polarity and cytoskeletal dynamics (Figure 2A), given they are highly interconnected developmental processes ^54^, associated with loss of trabeculation in mice and zebrafish ^1, 55, 56^. Accordingly, key genes required for cytoskeletal actin remodeling (*Arf4, Dock2, Sh3bp1, Arf6*), cytoskeletal organization (*Rac1, Rac2*), cell polarity (*Amotl1, Pard3, Nek3, Dlg1, Sipa1l3, Stk11, Wnt11*), cell division orientation (*Sapcd2*), motility (*Kif26b, Actr2*), and Fgf-ERK signaling (*Spry2, Fgf10*) were deregulated in *Nrg1^flox^*;*Tie2^Cre^* hearts (Figure 2A).

**Figure 2.**
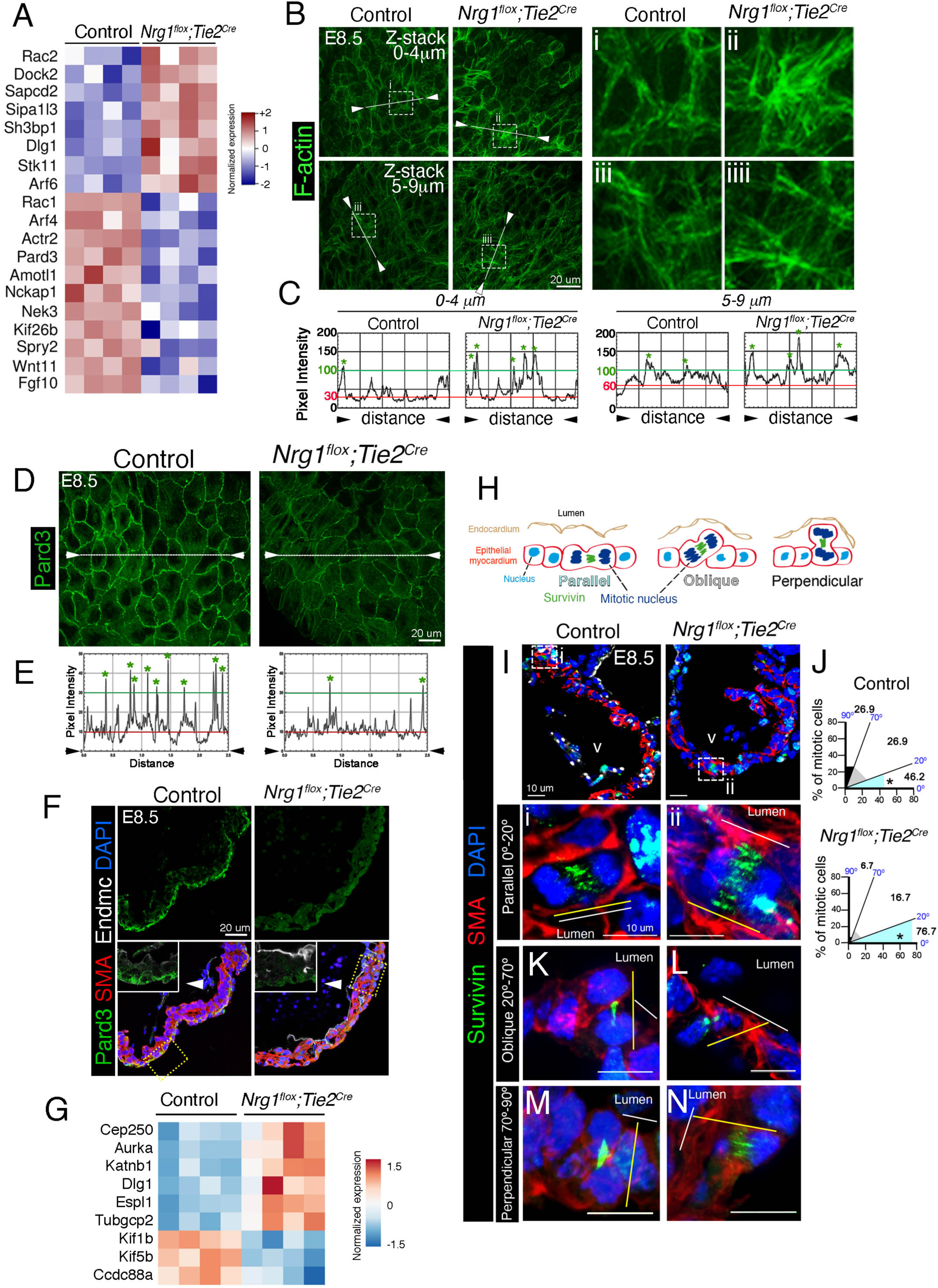
Nrg1 is required for polarity gene expression and cytoskeletal organization during trabeculation. (A) Enrichment analyses performed with Panther with the set of 1,219 differentially expressed genes in a *Nrg1^flox^;Tie2^Cre^*vs control comparison, against the Biological Process GO term database, identified “Establishment or maintenance of cell polarity” (GO:0007163) as enriched, with *P* value=0.0064.. The heatmap represents relative expression values for differentially expressed genes annotated with this GO term. **(B)** Whole mount phalloidin staining Z-stack projections of apical regions (from 0 to 4 μm) and baso-lateral regions (5 to 9 μm) of outer compact myocardium in E8.5 control and *Nrg1^flox^;Tie2^Cre^* heart sections. **(C)** Quantification of pixel intensity along a 10-pixel-wide trace (white lines in B). Red lines indicate that background intensities in control and *Nrg1^flox^;Tie2^Cre^*heart sections are similar. Green asterisks indicate values above gray intensity=100 (green lines). **(D)** Whole mount immunofluoresence of Pard3 in E8.5 control and *Nrg1^flox^;Tie2^Cre^*heart sections. **(E)** Quantification of pixel intensity along a 10-pixel-wide trace (white lines between arrowheads). Red lines indicate that background intensities in control and *Nrg1^flox^;Tie2^Cre^*heart sections are similar. Green asterisks indicate values above gray intensity=30 (green lines). **(F)** Pard3 immunodetection in transverse sections of E8.5 control and *Nrg1^flox^;Tie2^Cre^* embryos (green). Arrowheads to insets mark the apical domain of the compact myocardium. SMA, smooth muscle actin; Endmc, endomucin (white). DAPI counterstaining is shown in blue. **(G)** Heatmap of DEGs for components and regulators of the mitotic spindle in E9.5 control vs. *Nrg1^flox^;Tie2^Cre^*heart expression profiles. The color code represents normalized gene expression from –1.5 to +1.5. **(H)** Schematic of the analysis of mitotic-spindle orientation relative to the cardiac lumen: parallel (0° – 20°), oblique (20° – 70°), or perpendicular (70° – 90°). **(I)** Representative immunofluorescence images of mitotic cells stained against Survivin (green) and SMA (red) in control and *Nrg1^flox^;Tie2^Cre^* heart sections at E8.5. DAPI counterstaining is shown in blue. A general view of the E8.5 hearts is shown in the top panels. The white lines below indicate the reference plane of the cardiac lumen. The yellow lines indicate the orientation of the mitotic spindle. **(J)** Quantification of OCD in E8.5 control (N=3 embryos; n=26 mitotic figures) and *Nrg1^flox^;Tie2^Cre^* hearts (N=3 embryos; n=30 mitotic figures). *P* values were obtained by Fisher’s exact test. \**P*-value < 0.05. Scale bars, 20 μm (B, D, F); 15 μm (D); 10 μm (I).

The altered expression of actin remodeling genes in *Nrg1^flox^*;*Tie2^Cre^* mutants suggested the possibility of disrupted cardiomyocyte actin cytoarchitecture. Phalloidin staining of filamentous-actin (F-actin) in E8.5 control embryos revealed a uniform circumferential distribution of actin filaments attached to the plasma membrane (Figure 2B,C, Movies S3,S5). In contrast, in *Nrg1^flox^*;*Tie2^Cre^* hearts F-actin was non-uniformly distributed and appeared to concentrate in focal points at cell-cell junctions (Figure 2B,C, Movies S4,S6), suggesting that Nrg1 is required for proper arrangement of the actin cortical network. Consistent with this actin cytoskeletal defect, *Nrg1^flox^*;*Tie2^Cre^*mutants showed aberrant differential expression of genes sets for “Cytoskeleton” (GO:0005856) and “Regulation of actin polymerization” (GO: 0008064)(Figure S6A,B).

*Nrg1^flox^*;*Tie2^Cre^* hearts showed reduced transcription of the key cell polarity regulator *Pard3* (Figure 2A, Table S2, sheet 2). *Pard3* inactivation leads to impaired apico-basal (A-B) polarity in epithelial cells ^57^ and planar polarity defects in endothelial cells ^58^. “En face” examination of immunofluorescence (IF) staining in E8.5 control hearts revealed evenly distributed Pard3 protein at the apical/abluminal pole of ventricular cardiomyocytes (Figure 2D left, E). In contrast, in *Nrg1^flox^*;*Tie2^Cre^* hearts, Pard3 expression was strongly reduced (Figure 2D right, E). IF in heart sections confirmed Pard3 apical/abluminal pole localization in control ventricular cardiomyocytes at E8.5 (Figure 2F), a distribution that was maintained at E9.5 in the compact layer, whereas trabecular cardiomyocytes did not express Pard3 (Figure S7A,C,C’). This observation fits with the previous observations of loss of A-B polarity in cardiomyocytes that incorporate into nascent trabeculae ^1^. In contrast, Pard3 staining was severely depleted in E8.5 and E9.5 *Nrg1^flox^*;*Tie2^Cre^* ventricles (Figure 2F; Figure S7B,D,D’), potentially affecting A-B polarity of mutant cardiomyocytes. A second polarity complex, Crumbs2, has been shown to be required for the establishment of cardiomyocyte A-B polarity in zebrafish ^59^. We readily detected Crumbs2 at the apical pole of the compact layer in E8.5 control cardiomyocytes (Figure S7E,F), but this expression was strongly depleted in *Nrg1^flox^*;*Tie2^Cre^* mutants (Figure S7G,H), consistent with a potential loss of polarity in *Nrg1^flox^*;*Tie2^Cre^* cardiomyocytes. These data indicate that loss of Nrg1 signaling is associated with depleted expression of cell polarity proteins and a disorganized actin cytoskeletal network.

### Nrg1 is required for oriented division during trabeculation

Dynamic interplay between polarity complexes and the cytoskeleton dictates both the positioning of the mitotic spindle during mitosis and the orientation of cell division ^60^. Accordingly, several genes implicated in the regulation of mitotic spindle assembly were aberrantly expressed in *Nrg1^flox^*;*Tie2^Cre^*mutants, including *Cep250*, *Aurka*, *Katnb1*, *Dlg1*, *Tubgcp2*, *Kif1b* and *Kif5b*, among others (Figure 2G). Moreover, spindle orientation during trabeculation has been linked to the function and cellular distribution of adhesion complexes ^55^. The potential alteration in cell-cell adhesion molecules was manifested by differential regulation of “Cell adhesion” (GO: GO:0007155) and “Regulation of cell-matrix adhesion” (GO: 0001952) gene sets (Figure S6C,D). Overall, these data support the requirement for Nrg1-ErbB2-pERK signaling in the transcriptional regulation of cell polarity, adhesion and spindle orientation during trabeculation^55^.

Studies in mice show that ventricular cardiomyocytes divide obliquely or perpendicular to the lumen to give rise to the nascent trabeculae, whereas those that divide in parallel contribute to the outer compact myocardium ^6, 56^ (Figure 2H). We performed IF staining against Survivin to visualize the mitotic cleavage furrow and division axis during telophase (Figure 2I,J; Figure S7I-O). In E8.5 control hearts, 46.2% of mitotic cardiomyocyte divisions were parallel (between 0-20°) to the ventricular lumen (Figure 2I,i,J), whereas 26.9% were oblique (between 20-70°) with respect to the lumen (Figure 2I,J), and 26.9% perpendicular (between 70-90°; (Figure 2I,J). In contrast, in somite-matched E8.5 *Nrg1^flox^*;*Tie2^Cre^* mutants, parallel divisions were significantly more frequent (76.7%; Figure 2I,ii,J) and strong reductions were observed in oblique divisions (16.7%; Figure 2J,L) and perpendicular divisions (6.7%; Figure 2J,M,N). In E9.5 control embryos, parallel divisions remained predominant (61.4%) compared with perpendicular divisions (14%) or oblique divisions (24.6%; Figure S7I,K,M,O). In E9.5 *Nrg1^flox^*;*Tie2-CRE* mutants, up to 83.6% of divisions were parallel and 14.5% were oblique (Figure S7J,L,O), with perpendicular divisions significantly reduced to 1.8% (Figure S7N,O). Cardiomyocyte OCD is thus disrupted in E8.5-E9.5 *Nrg1^flox^*;*Tie2^Cre^*hearts, with parallel cardiomyocyte divisions favored at the expense of perpendicular divisions. We speculate that biased parallel division in *Nrg1^flox^*;*Tie2^Cre^*mutants could offset the observed reduction of proliferation at E9.5 and lead to increased compact myocardium thickness.

### Nrg1 is required for ventricular wall formation but not coronary vessels development

*Nrg1^flox^*;*Tie2^Cre^* embryos died between E11.5-E12.5 (Table S1, sheet 1), precluding analysis of late trabecular morphogenesis. To bypass this lethality, we bred *Nrg1^flox^* and *Cdh5^CreERT^*^2^ mice ^61^ to obtain mice with inducible pan-endothelial *Nrg1* deletion. Trabecular and compact wall growth was drastically hindered in E13.5 *Nrg1^flox^;Cdh5^CreERT^*^2^ mice after consecutive 4-Hydroxytamoxifen (4-OHT) administration at E10.5 and E11.5 (Figure S8A,B). At E16.5, this phenotype manifested as thinner rudimentary trabeculae and a significantly reduced ventricular wall thickness (Figure 3A-C). ISH analysis of early cardiac conduction system marker *Gja5*, the trabecular marker *Bmp10* and the compact myocardium marker *Hey2* revealed loss of trabeculae patterning and expansion of compact myocardium in *Nrg1^flox^;Cdh5^CreERT^*^2^ mutants (Figure 3D; Figure S8C-F). *Nrg1^flox^;Cdh5^CreERT^*^2^ mice survived until E18.5 (Table S1, sheet 3). Later 4-OHT induction, at E12.5 and E13.5, did not result in any obvious structural ventricular abnormalities at E18.5 (Figure S8G-K), suggesting that Nrg1–pErk signaling requirement precedes E13.5.

**Figure 3.**
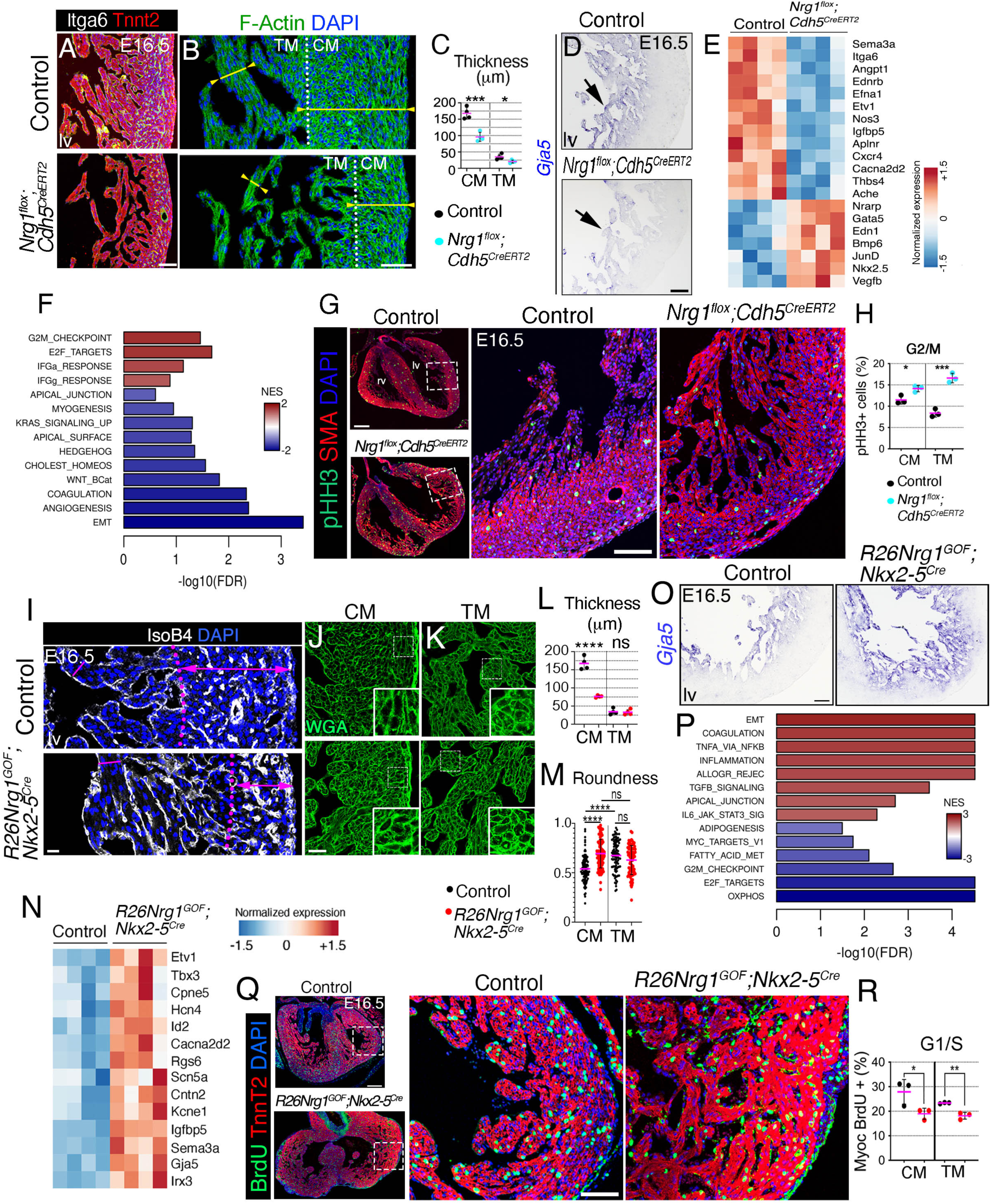
Nrg1 is required for trabecular growth, inner ventricular wall formation and its ectopic expression blocks ventricular compaction. (A) Immunofluorescence against integrin α6 (Itga6) and Troponin T2 (Tnnt2) in control and *Nrg1^flox^;Cdh5^CreERT^*^2^ heart transverse sections at E16.5. **(B)** Phalloidin (F-Actin) staining in E16.5 control and *Nrg1^flox^;Cdh5^CreERT^*^2^ heart sections. Dotted lines mark the separation between compact and trabecular myocardium. CM, compact myocardium; TM, trabecular myocardium. **(C)** Quantification of compact and trabecular myocardial thickness (n ≥3). **(D)** ISH of *Gja5* in E16.5 control and *Nrg1^flox^;Cdh5^CreERT^*^2^ embryos. Arrows mark trabecular myocardium. lv, left ventricle. **(E)** Heatmap of representative DEGs in E15.5 control vs. *Nrg1^flox^;Cdh5^CreERT^*^2^ heart expression profiles. The color code represents normalized gene expression from –1.5 to +1.5. **(F)** GSEA for E15.5 *Nrg1^flox^;Cdh5^CreERT^*^2^ vs. Control, against Hallmark gene sets. The bar plot represents enrichment data for 14 Hallmark gene sets at FDR qval < 0.25. Of these, 9 had FDR qval < 0.05. Scale bar indicates normalized enrichment score (NES) from –2 to +2. Positive (red) and negative (blue) scores, indicating enrichment or depletion, were found for 4 and 10 gene sets, respectively. Relative expression values for the “leading edge” genes associated to gene sets EMT and G2/M CHECKPOINT are presented as heatmaps in Figure S12. **(G)** left, phospho-histone 3 (pHH3)–positive immunofluorescence on sections of E16.5 control and *Nrg1^flox^;Cdh5^CreERT^*^2^ hearts (n=3 per genotype); right, boxed regions magnified **(H)** Quantification of % pHH3-positive myocardial cells. **(I)** DAPI staining in transverse sections of E16.5 control and *R26Nrg1^GOF^;Nkx2-5^Cre^*hearts. Double-headed arrows in the high-magnification views indicate myocardium thickness. Dotted lines demark the separation between CM and trabecular myocardium TM. **(J,K)** Wheat-germ-agglutinin (WGA)-FITC staining on transverse sections of CM and TM in E16.5 control and *R26Nrg1^GOF^;Nkx2-5^Cre^*hearts. **(L)** Quantification of CM and TM thickness in E16.5 control and *R26Nrg1^GOF^;Nkx2-5^Cre^* hearts (n=3 per genotype). **(M)** Quantification of CM and TM cell roundness in E16.5 control and *R26Nrg1^GOF^;Nkx2-5^Cre^*hearts sections (n=3 per genotype). **(N)** Heatmap of representative DEGs in E15.5 *R26Nrg1^GOF^;Nkx2-5^Cre^* vs. control heart expression profiles. The color code represents normalized gene expression from –1.5 to +1.5. **(O)** ISH for *Gja5* in E16.5 control and *R26Nrg1^GOF^;Nkx2-5^Cre^* heart sections. **(P)** GSEA for E15.5 *R26Nrg1^GOF^;Nkx2-5^Cre^* vs. control, against Hallmark gene sets. The bar plot represents enrichment data for a selection of 14 gene sets at FDR qval < 0.05. Scale bar indicates normalized enrichment score (NES) from –3 to +3. Positive (red) and negative (blue) scores, indicating enrichment or depletion, were found for 8 and 6 gene sets, respectively. **(Q)** left, BrdU-positive immunofluorescence on sections of E16.5 control and *R26Nrg1^GOF^;Nkx2-5^Cre^* hearts (n=3 per genotype); right, boxed regions magnified. **(R)** Quantification of % BrdU incorporation in CM and TM in control and *R26Nrg1^GOF^;Nkx2-5^Cre^*hearts (n=3 per genotype). \**P*-value < 0.05, ** *P*< 0.01, *** *P* < 0.001, **** *P* < 0.0001, ns=non-significant by Student t-test. Scale bars, 100 μm in A,D,O; 50 μm in B; 200 μm in G, 50 μm in G inset; 20 μm in I-K; 200 μm in Q, 50 μm in inset.

To determine the molecular basis of this phenotype, we performed an RNA-seq analysis on E15.5 control and *Nrg1^flox^;Cdh5^CreERT^*^2^ hearts (Table S3, sheet 1). Differential expression analysis identified 44 upregulated and 31 downregulated genes (B-H p-value<0.05) (Table S3, sheet 2). Differentially expressed genes were associated with pro-angiogenic and migratory processes (*Cxcr4, Vegfb*, *Aplnr1*, *Edn1, Ednrb*, and *Angpt1* ^62–65^; Figure 3E) and the development of the ventricular conduction system (VCS; *Sema3a*, *Efna1, Nkx2-5, Etv1,* and *Ache* ^66–70^; Figure 3E). Given that the endocardium gives rise to the ventricular inner wall vessels ^12, 14, 71^ and that trabeculae differentiate into the cardiac Purkinje Fiber network ^10^, these data raise the possibility that loss of endothelial Nrg1 expression from E11.5 onwards impairs the development of both the coronary arteries and the VCS.

In mice, the coronary arteries are located deep within the myocardial wall, whereas the veins are located superficially, beneath the epicardial layer ^72^. To assess the status of coronary vessel development in *Nrg1^flox^;Cdh5^CreERT^*^2^ mutants, we visualized the coronary network by whole-mount IF for IB4 (Figure S9). This analysis revealed that the organization and complexity of the coronary arterial tree was not obviously affected in E16.5 *Nrg1^flox^;Cdh5^CreERT^*^2^ mutant hearts (Figure S9A,B,D). Likewise, IF analysis of the venous endothelial marker Endomucin also revealed no appreciable disturbance of venous tree organization in *Nrg1^flox^;Cdh5^CreERT^*^2^ hearts (Figure S9E,F,H). Similar results were obtained with the vascular endothelial (but not endocardial)-specific *Pdgf^iCreERT^*^2^ Cre driver line (^73^; Figure S9C,D,G,H).

We next examined the expression of classical makers of the coronary veins (*CoupTFII*), arteries (*Dll4*), coronaries (*Fapbp4*), and endothelium (*Cdh5*) in developing heart by ISH, but again, did not find any obvious differences in the expression nor the distribution of these specific markers between E16.5 *Nrg1^flox^;Cdh5^CreERT^*^2^ mutants and control littermates (Figure S9I-P). To quantify the relative number of endothelial cells in the compact myocardium of E16.5 hearts we measured the percentage of IsolectinB4-positive (IB4 +) signal in the compact myocardium (Figure S10A-C). Again, there were no differences between *Nrg1^flox^;Cdh5^CreERT^*^2^ and control hearts (Figure S10C), consistent with absence of coronary vascular defect in Nrg1 mutants. Finally, we used SM22 and Notch3 immunostainings to determine the extent of coronary vessel smooth muscle cell differentiation and pericyte coverage, respectively ^72^. Consistent with the absence of coronary defects, these parameters were also comparable between *Nrg1^flox^;Cdh5^CreERT^*^2^ and control hearts (Figure S10D,E).

These data indicate that lack of an inner compact wall does not appreciably impact the development of the coronary tree in *Nrg1^flox^;Cdh5^CreERT^*^2^ hearts. A reasonable interpretation is that the potential below-normal endocardial contribution to coronary vasculogenesis in these mutants could be balanced by compensatory mechanisms involving alternative sources of endothelial cells^74^.

### Nrg1 regulates motility and G2/M progression during ventricular wall growth

To identify mechanisms underlying the *Nrg1^flox^;Cdh5^CreERT^*^2^ phenotype, we performed GSEA. Among the “HALLMARK” gene sets, the most depleted pathway was EMT (-log(FDR) p-value < 0.05; Figure 3F; Table S3, sheet 3). The subset of leading-edge depleted genes was associated with cell cytoskeleton, ECM composition, modifiers of the ECM, and transmembrane and secreted proteins including adhesion molecules (Figure S11A,B). Other depleted gene sets evoked cell shape and polarity, (APICAL_SURFACE, APICAL_JUNCTION), and relevant pro-migratory signaling pathways, (WNT_BETA_CATENIN_SIGNALING, HEDGEHOG_SIGNALING) (Figure 3F).

Furthermore, depleted CHOLESTEROL_HOMEOSTASIS was consistent with the essential role of Nrg1-ErbB4 signaling in cholesterol metabolism ^75^. In contrast, several over-represented pathways (G2/M_CHECKPOINT, E2F_TARGETS) suggested a proliferative defect (Figure 3F). The “leading-edge” genes associated with these pathways were linked to the cell cycle, chromatin, DNA damage response, DNA repair, DNA replication, and mitosis (Figure S11C), whereas nuclear transport, transcription regulation pathways and, importantly, the cytoskeleton (Figure S11B,C), emphasize the nucleus-cytoskeleton connection.

To support this finding, we examined E16.5 *Nrg1^flox^;Cdh5^CreERT^*^2^ mutants for possible alterations to the G1/S and G2/M cell-cycle phases by respectively measuring BrdU incorporation and positivity for the early mitosis marker phospho-histone H3 (pHH3) in cardiomyocytes. Compared with stage-matched control hearts, E16.5 *Nrg1^flox^;Cdh5^CreERT^*^2^ hearts contained a significantly higher proportion of pHH3-positive (pHH3+) cells, especially in trabeculae (Figure 3G,H). These differences were already manifest at E12.5 (Figure S12A,B,R). There was no alteration in the proportion of BrdU+ cells in compact or trabecular myocardium, either at E13.5 (Figure S12C,D,G) or at E16.5 (Figure S12E,F,G). The absence of effect of Nrg1 depletion on proliferation at either E12.5 or E16.5 was confirmed by staining against Ki67 antigen, which preferentially marks the S-G2 phases (Figure S12H-K,R). These data suggest that cell-cycle progression through the G2/M phase is delayed or prolonged in *Nrg1* mutants, while the G1/S phase is unaffected.

To further characterize cardiomyocyte growth retardation in the *Nrg1^flox^;Cdh5^CreERT^*^2^ mutants, we examined p27^Kip1^ protein expression, as both an effector of pERK ^76^ and a regulator of cell cycle progression at both G1/S ^77^ and G2/M ^78^ phases. p27 expression was readily increased in *Nrg1^flox^;Cdh5^CreERT^*^2^ compact and trabecular myocardium at E12.5 (Figure S12L,M,R), correlating with the increase in pHH3 and a potentially delayed G2/M phase. This difference persisted in compact myocardium at E16.5, although the overall p27 staining in both control and mutant was severely attenuated by this timepoint (Figure S12N-Q,R).

Overall, the analysis of the *Nrg1^flox^;Cdh5^CreERT^*^2^ transcriptome suggests that Nrg1 regulates trabecular growth and chamber wall formation by coordinating cell migration and G2/M phase progression.

### Ectopic Nrg1 expression induces trabeculation of ventricular wall myocardium

To gain further insight into the role of Nrg1 in chamber development, we created a conditional Nrg1 gain-of-function model. For this, we generated a transgenic line bearing a *Rosa26-floxNeoSTOPflox-Nrg1-EGFP* expression cassette (Figure S13A,B; see *METHODS*). Nkx2-5^Cre^-mediated removal of the NeoSTOP sequence resulted in the expression of both the GFP reporter (Figure S13C,D) and Nrg1 protein (Figure S13E,F). Examination of E16.5 embryos showed that *R26Nrg1^GOF^;Nkx2-5^Cre^*transgenic hearts were developmentally arrested, with an appearance similar to that of E12.5 wild-type hearts (Figure S14A,B). This developmental arrest was further evidenced by the presence of a double-outlet right ventricle, leading to overriding aorta (Figure S14A) and a ventricular septal defect (muscular and peri-membranous; Figure S14B). E16.5 *R26Nrg1^GOF^;Nkx2-5^Cre^*transgenic hearts also displayed superficial outgrowths resembling fistulae (Figure S14Ci,ii), which were possibly manifestations of direct communication between the endocardium and epicardium caused by considerable thinning of the compact myocardium (Figure 3I,L). *R26Nrg1^GOF^;Nkx2-5^Cre^*mice die by E18.5 (Table S1, sheet 4).

Histological analysis of E16.5 hearts showed that the transgenic epicardium consisted of 3–4 cell layers instead of a single cell layer (Figure S14D), and the sub-epicardial layer was extensively infiltrated by endothelial cells but largely devoid of coherent large vessel conduits (Figure 3I; Figure S14D). Cardiomyocyte morphology (roundness) in the outer compact myocardium of *R26Nrg1^GOF^;Nkx2-5^Cre^*transgenic hearts resembled more the trabecular myocardium of controls than the corresponding compact myocardium (Figure 3J,K,M). Moreover, expression of ITGα6 was expanded in E10.5 *R26Nrg1^GOF^;Nkx2-5^Cre^*hearts from the trabeculae to the compact layer (Figure S14E,F). Thus, Nrg1 overexpression causes the compact myocardium to become trabecular-like.

To define the molecular underpinnings of this phenotype and to compare with the one obtained from *Nrg1^flox^;Cdh5^CreERT^*^2^ hearts, we performed RNA-seq on E15.5 transgenic hearts. Differential expression analysis identified > 4,500 DEGs in *R26Nrg1^GOF^;Nkx2-5^Cre^*hearts (FDR q-value<0.05; Table S4, sheets 1 and 2). As expected, the expression of a subset of His-Purkinje-specific genes ^79^ was broadly upregulated (Figure 3N). ISH analysis revealed expansion of *Gja5* transcription towards the compact myocardium (Figure 3O), consistent with the proposed role of Nrg1 as a driver of VCS specification and differentiation ^80^. Pathway analysis revealed the enrichment for pro-migratory processes (EMT, APICAL_JUNCTION, KRAS_SIGNALING_UP) in *R26Nrg1^GOF^;Nkx2-5^Cre^*hearts (Figure 3P), involving expression of the EMT transcription factor drivers Snail and Twist genes (Figure S15A; Table S4, sheets 2 and 3), as well as several key EMT signaling pathways, including TGFβ (TGF_BETA_SIGNALING) and BMP (Figure 3P; Figure S15B,C; Table S4, sheets 2 and 3). Conversely, metabolic pathways (ie.: OXIDATIVE_PHOSPHORYLATION, FATTY_ACID_METABOLISM, ADIPOGENESIS, PROTEIN_SECRETION, MYOGENESIS, CHOLESTEROL_HOMEOSTASIS), associated with MYC activity (MYC_TARGET_V1, MYC_TARGET_V2) (Figure 3P, Table S4, sheets 2 and 3), were depleted, revealing an impairment of cardiac metabolic maturation. In support of this maturational defect, we examined the expression of intermediate filament genes as they have been involved in lineage determination and organ maturation ^81^. Several genes encoding lamins, nestin and various keratin proteins were differentially regulated in *R26Nrg1^GOF^;Nkx2-5^Cre^* hearts (Figure S15D). Notably, mis-expression of the intermediate filament protein Nestin, has been characterized as a marker of cardiac immaturity ^82^. Thus, in E16.5 control hearts, Nestin expression was weak in trabecular myocardium but strong in endothelium and epicardium (Figure S14Gi,i’). In contrast, in *R26Nrg1^GOF^;Nkx2-5^Cre^* transgenic hearts, Nestin expression was substantial in the trabecular layer and expanded sub-epicardium layer, devoid of α-smooth-muscle actin (SMA) staining (Figure S14Hii,ií). Overall, Nrg1 overexpression alters the relative proportion of constituent ventricular wall cell types and causes the compact myocardium to become trabecular-like

### Ectopic Nrg1 expression causes cell cycle arrest and a pro-senescence phenotype

The metabolic defect was also associated with the upregulation of cytotoxic stress (IL6_JAK_STAT3_SIGNALING, HYPOXIA, IL2_STAT5_SIGNALING, UNFOLDED PROTEIN RESPONSE), pro-inflammatory (INFLAMMATORY_RESPONSE, COAGULATION, COMPLEMENT, ALLOGRAFT_REJECTION, INTERFERON_GAMMA_RESPONSE), and pro-senescence (TNA_SIGNALING_VIA_NFKB) gene sets (Figure 3P; Figure S15E,F; Table S4, sheet 3). The activation of cellular stress pathways suppresses mitogenic responses and promotes senescence ^83^, and have been previously associated with the so-called “hypermitotic” cell cycle arrest phenomenon ^84^.

Indeed, the depletion of cell-cycle–related gene sets (E2F_TARGETS and G2M_CHECKPOINT) (Figure 3P), suggested a proliferation defect associated with the fetal developmental arrest of *R26Nrg1^GOF^;Nkx2-5^Cre^*embryos. We examined E12.5 and E16.5 *R26Nrg1^GOF^;Nkx2-5^Cre^*mutants for possible alterations to the G1/S and G2/M cell-cycle phases. The proportion of BrdU+ cells was lower in the compact and trabecular myocardium of E16.5 *R26Nrg1^GOF^;Nkx2-5^Cre^* hearts (Figure 3Q,R), suggesting either decreased proliferation rate, or delayed G1/S progression. The inhibitory effect of Nrg1 expression on compact and trabecular cardiomyocytes proliferation was confirmed by staining against Ki67 antigen (Figure S16A-D). In contrast, the pHH3+ cell indexes in compact myocardium were similar in E12.5 and E16.5 control and *R26Nrg1^GOF^;Nkx2-5^Cre^* transgenic hearts (Figure S16E,F,I), although in trabecular myocardium at E12.5, pHH3+ staining was increased, but no longer at E16.5 (Figure S16E,F,I). These data suggest that the proliferation rate across the ventricular wall is homogeneously decreased in *R26Nrg1^GOF^;Nkx2-5^Cre^* transgenic hearts, consistent with the suggested myocardial “trabeculation” of the compact layer.

Closer inspection of the E2F_TARGET genes revealed several key cell cycle inhibitors that were either increased (*cdkn1a*/p21) or decreased (*cdkn1b*/p27*, cdkn2c*/p18) (Figure S15G), as potentially underlying the growth retardation defect. We examined the expression of the p27 inhibitor of cell cycle progression ^85^ in E12.5 *R26Nrg1^GOF^;Nkx2-5^Cre^* embryos, and found it to be markedly increased in compact and trabecular myocardium compared to controls (Figure S16G,I), correlating with the decrease in BrdU+ labelling (Figure 3Q,R) and a potentially delayed G1/S phase. Here, *cdkn1b* gene expression did not correlate with its protein expression, likely because regulation of p27 is predominantly post-transcriptional ^86^. This increased p27 expression persisted in compact myocardium at E16.5, although the overall p27 staining in both control and mutant was severely attenuated by this stage (Figure S16H,I). Overall, these data indicate that Nrg1 overexpression promotes cell cycle arrest and a prosenescence phenotype.

### Nrg1 regulates ventricular patterning mediated by cytoskeletal dynamics

Analysis of chamber trabecular and compact myocardium markers revealed profoundly altered myocardial wall structure and patterning in both the *Nrg1^flox^;Cdh5^CreERT^*^2^ mutant and *R26Nrg1^GOF^;Nkx2-5^Cre^* transgenic hearts (Figure 4A-L). At E16.5, SMA is normally confined to the immature and proliferative compact myocardium (^87^; Figure 4D) but extended along the length of trabeculae in *Nrg1^flox^;Cdh5^CreERT^*^2^ hearts (Figure 4E), consistent with defective trabecular maturation and maintenance of compact myocardium identity. In contrast, SMA was markedly reduced in the compact myocardium of *R26Nrg1^GOF^;Nkx2-5^Cre^*transgenic hearts (Figure 4F), in line with a loss of compact myocardium identity and gain of trabecular one. In agreement with these findings, ISH showed that *Hey2* expression abnormally extended into the trabeculae of *Nrg1^flox^;Cdh5^CreERT^*^2^ hearts, contrasting its normal restriction to the compact myocardium (Figure 4G,H), and was almost completely absent from *R26Nrg1^GOF^;Nkx2-5^Cre^* ventricles (Figure 4G,I), reflecting loss of compact myocardium identity. Conversely, *Bmp10* expression was attenuated in the trabeculae of *Nrg1^flox^;Cdh5^CreERT^*^2^ mutants (Figure 4J,K) and extended into the compact layer in *R26Nrg1^GOF^;Nkx2-5^Cre^* transgenic embryos (Figure 4J,L), reflecting acquisition of trabecular identity. Thus, Nrg1 loss of function causes trabecular myocardium to become compact-like, whereas Nrg1 gain-of-function causes compact myocardium to become trabecular-like.

**Figure 4.**
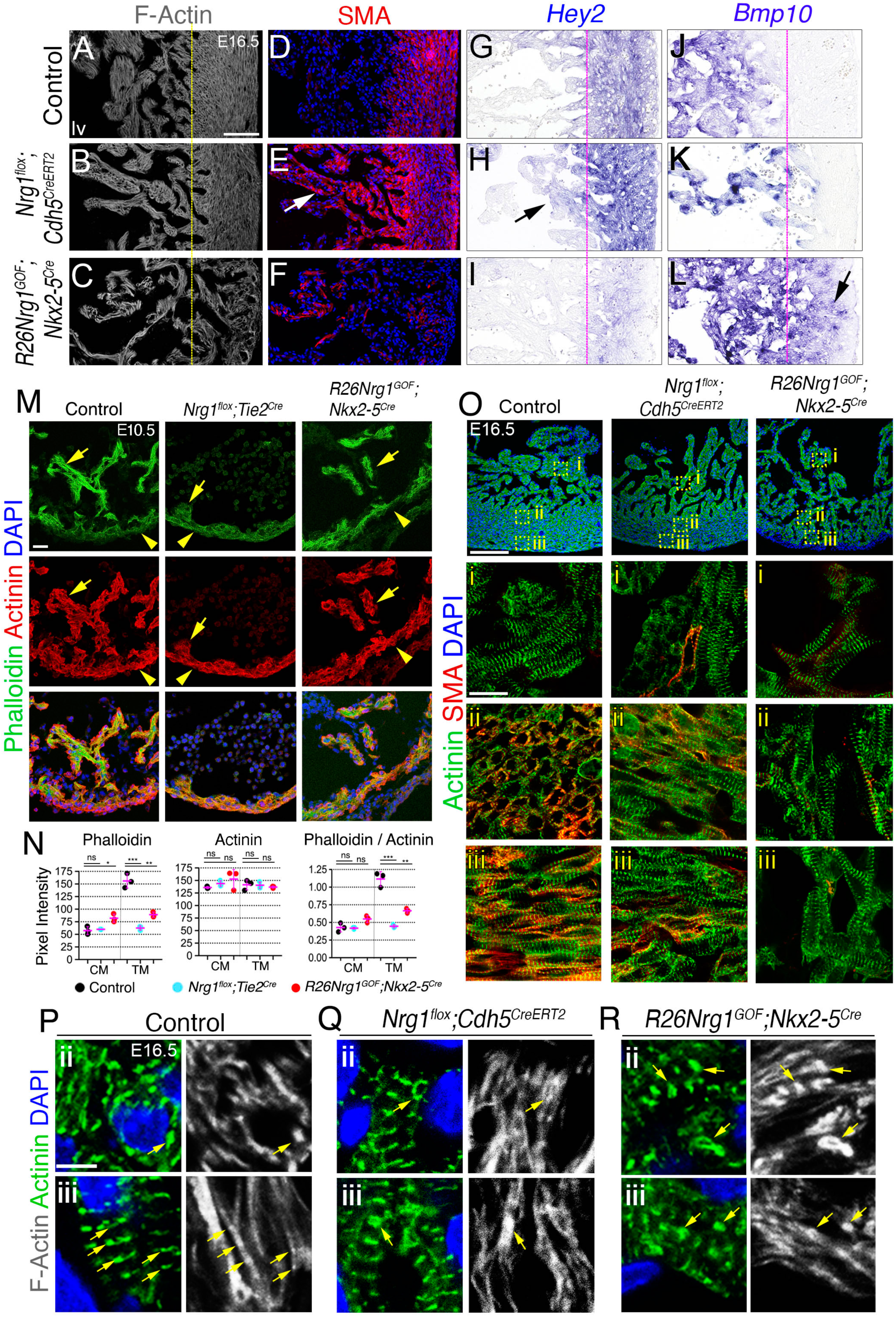
Ventricular patterning is mediated by cytoskeletal dynamics. (A-F) Phalloidin (F-actin) staining and immunodetection of smooth muscle actin (SMA) in E16.5 left ventricular sections from control **(A, D)**, *Nrg1^flox^;Cdh5^CreERT^*^2^ **(B, E)**, and *R26Nrg1^GOF^;Nkx2-5^Cre^* hearts **(C, F)**. Yellow lines mark the position of the morphological border between trabecular myocardium (TM) and compact myocardium (CM) in control heart sections, showing how this border shifts in *Nrg1^flox^;Cdh5^CreERT^*^2^ and *R26Nrg1^GOF^;Nkx2-5^Cre^* hearts. The arrow marks strong SMA immunostaining in TM relative to CM in the *Nrg1^flox^;Cdh5^Cre^* heart. Note also the reduced SMA expression in CM of *R26Nrg1^GOF^;Nkx2-5^Cre^*hearts **(F). (G-L)** ISH for *Hey2* and *BMP10* in E16.5 left ventricular sections from control **(G, J)**, *Nrg1^flox^;Cdh5^CreERT^*^2^ **(H, K)**, and *R26Nrg1^GOF^;Nkx2-5-Cre* hearts **(I, L)**. Pink lines mark the position of the morphological border between TM and CM in control heart sections, showing how this border shifts in *Nrg1^flox^;Cdh5^CreERT^*^2^ and *R26Nrg1^GOF^;Nkx2-5^Cre^* hearts. Arrows mark ectopic expression of *Hey2* and *Bmp10* in *Nrg1^flox^;Cdh5^CreERT^*^2^ and *R26Nrg1^GOF^;Nkx2-5^Cre^* hearts, respectively. Note also the reduced *Hey2* expression in CM of *R26Nrg1^GOF^;Nkx2-5^Cre^*hearts. (**M**) F-actin (Phalloidin) and α-actinin immunodetection in E10.5 control, *Nrg1^flox^;Tie2^Cre^*and *R26Nrg1^GOF^;Nkx2-5^Cre^*hearts. Yellow arrows indicate TM and arrowheads CM. **(N)** Quantification of Phalloidin and α-actinin immunofluorescence, and Phalloidin/α-actinin ratios in heart sections from E10.5 control, *Nrg1^flox^;Cdh5^CreERT^*^2^ and *R26Nrg1^GOF^;Nkx2-5^Cre^* embryos. **(O)** α-actinin and α-SMA immunodetection in E16.5 control, *Nrg1^flox^;Cdh5^CreERT^*^2^ and *R26Nrg1^GOF^;Nkx2-5^Cre^* hearts. Panels i, ii, and iii show high-magnification views of TM, inner CM, and outer CM, respectively. **(P-R)** Detail of F-actin and α-actinin double immunofluorescence at E16.5 in inner CM (ii) and outer CM (iii). Yellow arrows mark Z-bands and highlight aberrant Z-band structures in *Nrg1^flox^;Cdh5^CreERT^*^2^ and *R26Nrg1^GOF^;Nkx2-5^Cre^*hearts. Scale bars, 100 μm in A-I,O; 20 μm in M; 10μm in O, i-iii; 5μm in P-R.

Tissue patterning is determined, in part, by the organization of cytoskeletal components ^88^. In cardiomyocytes, these components form the sarcomere, which are bundles of polymerized actin crosslinked by α-actinin (Z-line) and arranged in a stacked striated pattern throughout muscle tissue. We therefore examined changes in the actin cytoskeleton potentially underlying the patterning defects, by examining F-actin (Phalloidin) and α-actinin (Figure 4M-O). In control E10.5 hearts, the intensity of Phalloidin staining was markedly higher in trabecular compared to compact myocardium, in contrast to α-actinin, which displayed similar intensity of staining (Figure 4M,N). Phalloidin staining in trabecular myocardium was normalized to compact myocardium intensity in E10.5 *Nrg1^flox^;Tie2^Cre^* mutants, while staining in compact myocardium was normalized to trabecular myocardium level in E10.5 *R26Nrg1^GOF^;Nkx2-5^Cre^* hearts (Figure 4M,N). Thus, the contrast between F-actin staining intensities was absent in both *Nrg1^flox^;Tie2^Cre^* and *R26Nrg1^GOF^;Nkx2-5^Cre^*hearts, which is consistent with the loss of trabecular and compact myocardium identities, respectively. We investigated the arrangement of cardiomyocytes by visualizing their sarcomeres using IF against α-actinin (Figure 4O). The sarcomeres in trabecular cardiomyocytes of E16.5 control, *Nrg1^flox^;Cdh5^CreERT^*^2^ and *R26Nrg1^GOF^;Nkx2-5^Cre^* hearts exhibited a well organized striation pattern (Figure 4Oi). In the compact myocardium of control hearts, cardiomyocytes displayed two distinct fiber patterns based on their striation; those in the inner compact myocardium exhibited a circular striation pattern (Figure 4Oii), while cardiomyocytes in the outer compact myocardium displayed a similar fiber orientation as in the trabeculae (Figure 4Oiii). Analysis of sarcomere fiber orientation revealed that *Nrg1^flox^;Cdh5^CreERT^*^2^ hearts exhibited a deficiency in the inner compact myocardium layer, while maintaining a normal outer compact myocardium (Figure 4Oii, iii). Conversely, the compact myocardium of *R26Nrg1^GOF^;Nkx2-5^Cre^* ventricles displayed a sarcomere fiber orientation akin to that of the trabecular myocardium (Figure 4Oi-iii). Upon closer inspection, *Nrg1^flox^;Cdh5^CreERT^*^2^ compact-myocardium sarcomeres appeared slightly more diffuse than their control counterparts and contained ring-like structures (Figure 4P,Qii,iii), suggesting disruption of Z-line alignment. The striated sarcomere structure revealed by α-actinin was also broadly evident from F-actin staining, including the presence of ring-like structures in *Nrg1^flox^;Cdh5^CreERT^*^2^ hearts (Figure 4Qii,iii), suggesting mis-arrangement of F-actin branching. Similar ring-like structures were detected in the compact myocardium sarcomeres of *R26Nrg1^GOF^;Nkx2-5^Cre^* hearts (Figure 4Rii,iii). Overall, these data indicate that above-normal or below-normal Nrg1 expression causing the ventricular patterning defects are associated with disrupted F-actin organization and sarcomere Z-line positioning.

### Nrg1 modulates pErk-dependent Yap1 S274 phosphorylation during trabeculation

Having identified myocardial pErk as a read-out of Nrg1 signaling during trabeculation (see Figure 1K-M; Figure S2J-N;), we monitored pErk expression in *R26Nrg1^GOF^;Nkx2-5^Cre^*transgenic hearts. In E9.5 control hearts, pErk was expressed in trabeculae and in cardiomyocytes close to the ventricular lumen (Figure 5A), and additionally in some epicardial cells at E11.5 (Figure 5C). In contrast, E9.5-E11.5 *R26Nrg1^GOF^;Nkx2-5^Cre^* transgenic hearts showed ectopic pErk expression in the compact myocardium and sub-epicardial layers (Figure 5B,D). From E13.5 onwards, pErk was no longer detected in the myocardium of control hearts (Figure S2M,N), but was expressed in coronary endothelial cells (Figure S17A), similarly to the observation in *Nrg1^flox^;Cadh5^CreERT^*^2^ hearts (Figure S17B). At E16.5, pErk was no longer expressed ectopically in *R26Nrg1^GOF^;Nkx2-5^Cre^*myocardium (Figure S17C).

**Figure 5.**
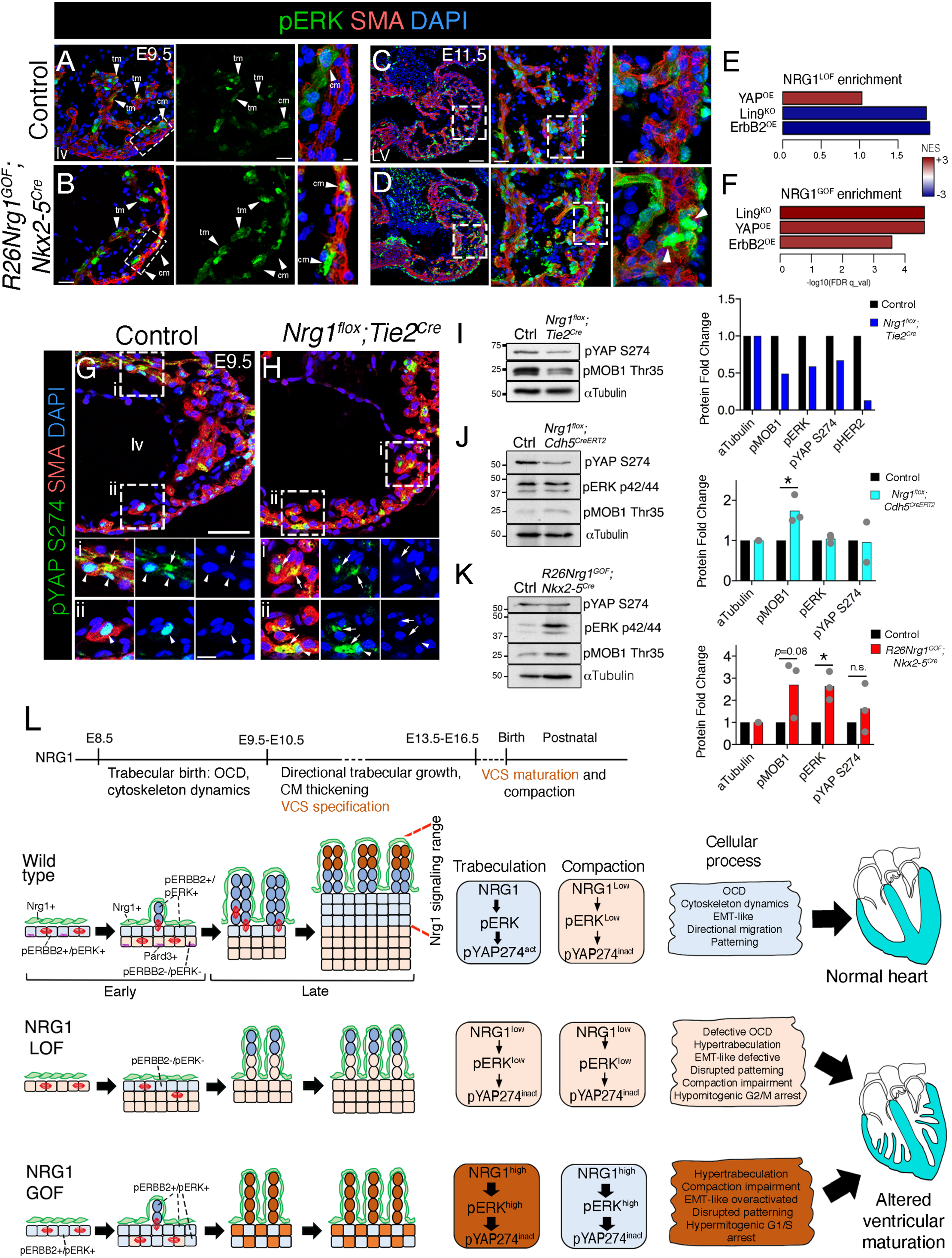
Ectopic pErk expression in *R26Nrg1^GOF^;Nkx2-5^Cre^* compact myocardium and Nrg1-dependent pYap1 S274 regulation during trabeculation. (A-C) Immunofluorescence for pERK in left ventricular sections from E9.5 **(A, B)** and E11.5 embryos **(C, D)**. Arrowheads mark labeled cardiomyocytes in trabecular (tm) and compact (cm) myocardium. **(E)** GSEA for E15.5 *Nrg1^flox^;Cdh5^CreERT^*^2^ vs. Control and **(F)** E15.5 *R26Nrg1^GOF^;Nkx2-5^Cre^*vs. Control, against gene sets consisting of collections of differentially expressed genes derived from ERBB2^OE^ vs. Control ^89^ and Lin9KO vs. Control^90^, or direct YAP1-target genes^92^. The barplots represent enrichment data at FDR qval < 0.25, and the color scale indicates normalized enrichment score (NES) from –3 to +3. **(G)** Immunofluorescence detection of pYAP S274 in E9.5 control and (H) *Nrg1^flox^;Tie2^Cre^*heart sections. Arrows mark cytoplasmic signal, arrowheads nuclear signal. **(I)** Western blot analysis and quantification of pYAP S274 and pMOB1 Thr35 in E9.5 control and *Nrg1^flox^;Tie2^Cre^* ventricles (pool of n=3 per genotype). **(J)** Western blot analysis of pYAP S274, pERK 42/44, and pMOB1 Thr35 in E16.5 control and *Nrg1^flox^;Cdh5^CreERT^*^2^, and **(K)** control and *R26Nrg1^GOF^;Nkx2-5^Cre^* ventricles (n=1 per genotype), and quantifications (n=2-3). \**P*-value < 0.05, ns=non-significant by Student t-testα−Tubulin was used as a gel-loading control and for normalization in quantifications. Scale bars, 100 μm, 20 μm, and 10 μm from low to high magnification view in A, B; 100 μm, 30 μm, and 10 μm in C, D; 50 μm in G,H; i,ii, 10 μm**. (L)** Nrg1 regulates cardiomyocyte dynamics, cell cycle progression, and maturation during ventricular chamber morphogenesis. Wild type (E8.5-E10.5), the early myocardium is characterized by parallel divisions in the outer myocardial layer and perpendicular cell divisions in the inner myocardial layer. Nrg1 released from the endocardium (cells in green) activates pErbB2 and pErk in the trabecular layer (cells in blue), promoting pYap S274 nuclear localization (pYap S274^act^). This signaling is associated with the activation of actin cytoskeleton, EMT-like dynamics, directional migration, and chamber patterning (compact vs. trabecular). Nrg1 LOF (E8.5-E10.5), deficient Nrg1 signaling leads to reduced perpendicular divisions in favor of parallel ones (defective OCD), causing compact myocardium thickening and trabeculae underdevelopment. These changes are associated with the depletion of apico-basal polarity markers, disorganized actin cytoskeleton, low pErk and pYap S274 inactivation (pYap S274^inact^). Wild-type (E10.5-E16.5), trabeculae elongate by directional migration from the proliferative outer compact myocardium (cells in beige) towards the inner compact myocardium (cells in blue). Nrg1 is required for specification of the ventricular conduction system (VCS) in trabeculae (brown cells) but not for compaction in late gestation or postnatally. LOF (E10.5-E16.5), Nrg1 depletion and low pErk, trabecular myocardium expresses compact myocardium genes, leading to rudimentary and mis-patterned trabeculae, blunted inner myocardial wall growth, and hypomitogenic cell cycle arrest. GOF (E10.5-E16.5), cardiac Nrg1 overexpression and high pErk, disrupts patterning by trabeculating the compact myocardium and impairing compaction. Hypermitogenic signaling causes cell cycle arrest in cardiomyocytes and onset of senescence. Both Nrg1 LOF and GOF lead to altered ventricular chamber maturation and cardiac developmental arrest.

To support these findings, we examined the expression of pErbB2 and pErbB4 during compaction. In E16.5 control hearts, pErbB2 and pErbB4 were both strongly expressed in coronary endothelium, sparsely expressed in endocardium, and absent from myocardium (Figure S17D,G). pErbB2 was also readily detected in epicardium (Figure S17D). In *Nrg1^flox^;Cadh5^CreERT^*^2^ hearts, the strong coronary endothelial expression and sparse endocardial expression were maintained (Figure S17E,H). In *R26Nrg1^GOF^;Nkx2-5^Cre^* hearts, strong pErbB2 and pErbB4 expression was detected in both endothelium and endocardium, and remained absent from myocardium (Figure S17F,I). Thus, during compaction, pErbB2 and pErbB4 are no longer expressed in myocardium, and cannot be induced by Nrg1 in this tissue.

We noticed that the *R26Nrg1^GOF^;Nkx2-5^Cre^*gene signature closely mirrored gene signatures found in adult mouse hearts overexpressing the ErbB2 receptor (ErbB2^OE^) ^89^ and in fetal hearts deficient for the activator MuvB core complex component Lin9 (Lin9^KO^) ^90^ (Figure S17J,K). Notably, the EMT and oxidative phosphorylation gene signatures were respectively enriched and depleted in the transcriptomes of all three of these models, whereas the opposite change occurred after Nrg1 deletion (Figure 3F, S17L). ErbB2-mediated cardiac regenerative phenotypes are mediated, in part, by Yap1 ^89^, and there is significant overlap between the conserved Yap1-regulated and Lin9-dependent gene signatures ^91^. Enrichment analysis revealed that many of the differentially-regulated genes in ErbB2^OE^ and Lin9^KO^ hearts were also significantly altered in the Nrg1 transcriptomes (Figure 5E,F; Figure S17L). Moreover, the E15.5 *R26Nrg1^GOF^;Nkx2-5^Cre^* transcriptome displayed significant overlap with the Yap1 target gene signature ^92^ (Figure S18A). Part of this enriched gene signature corresponded to the EMT pathway (Figure S17L), suggesting that a significant portion of the Nrg1 transcriptome involving cytoskeletal genes might also be YAP-dependent.

The activation/inactivation of Yap by canonical Hippo-dependent and non-Hippo dependent phosphorylations has been linked to mechano-transduction and mitosis during cardiac regeneration ^89^. ErbB2 signaling via Erk elicits cytoskeletal changes with an altered mechanical state and downstream YapS274 phosphorylation, resulting in pYap activation and nuclear localization ^89^. Given the enrichment of Yap1-target genes in *R26Nrg1^GOF^;Nkx2-5^Cre^* hearts, we examined the Nrg1 mutants for Hippo (Mob1)-dependent Yap1 phosphorylation on S112 (S127 in human) and pErk and mitosis-associated Yap1 phosphorylation on S274 (S289 in human) ^93, 94^. In control E9.5 hearts, pYap S112 was strongly expressed in the cytoplasm of endocardial cells and weakly expressed in some cardiomyocytes (Figure S18B). In *Nrg1^flox^;Tie2^Cre^*hearts, pYap S112, while maintained in endocardium, was not visible in cardiomyocytes (Figure S18C). Interestingly, pMob1-Thr35 expression was also below normal (Figure 5I), suggesting that Hippo-dependent Yap1 nuclear localization would be higher in E9.5 *Nrg1^flox^;Tie2^Cre^* hearts. However, at E13.5, pYap S112 was present exclusively in endocardial cells in both control and *Nrg1^flox^;Cadh5^CreERT^*^2^ hearts (Figure S18D,E), and the same restriction was observed at E16.5 in control and *R26Nrg1^GOF^;Nkx2-5^Cre^*hearts (Figure S18F,G). Thus, pYap S112 expression and Hippo-regulated Yap activity appear to be dependent on Nrg1 at E9.5, but independent of Nrg at E13.5 or E16.5.

At E9.5, pYap S274 was localized in the nuclei of myocardial cells in control hearts (Figure 5G) but was mostly cytoplasmic in *Nrg1^flox^;Tie2^Cre^* hearts (Figure 5H). In agreement with this, Western blot analysis revealed below-control pYap S274 expression in *Nrg1^flox^;Tie2^Cre^*hearts (Figure 5I). Consistent with the dependence of pYap S274 on Erk phosphorylation ^89^, pErk expression was also below control (Figure 1M). In contrast, at E16.5, pYap S274 IF labeled only scattered cells in the myocardium of control, *Nrg1^flox^;Cdh5^CreERT^*^2^, and *R26Nrg1^GOF^;Nkx2-5^Cre^* hearts (Figure S18H-J), although Western blot analysis detected above control-level expression of pMob1-Thr35 in both Nrg1 LOF and GOF mutants (Figure 5J,K). These data suggest that Erk phosphorylation, pYap S274 cellular distribution and Hippo/Mob1 activity in cardiomyocytes, are dependent on Nrg1 during trabeculation but not during compaction

## Discussion

The results presented here show that Nrg1 regulates a distribution of polarized morphologies and division dynamics across the ventricular wall (Figure 5L). We interpret our findings in light of a body of work showing that trabeculation is a dynamic process involving transient disruption of the integrity of the myocardial cell layer. In zebrafish this manifests as an EMT-like process that involves changes in the actin cytoskeleton network, resulting in actomyosin polarization, apical constriction, depolarization, and delamination without division ^1, 3–5, 53, 95^. In mice, the realignment of the dividing axis of cardiomyocytes undergoing trabeculation involves transient disruption and re-positioning of polarity, focal cell and matrix adhesion complexes ^6, 7, 56^. Furthermore, interlinked posttranscriptional and transcriptional cascades converge to drive dynamic changes in the contractile actomyosin network, influence force transmission, to control spindle orientation and directional migration ^54, 60^.

We infer from our transcriptomics data that Nrg1 regulates the expression of genes required for cell shape and morphology, including key polarity complex components, with depletion of the apical junction and polarity proteins Pard3 and Crumbs2, which have been linked to oriented division and trabeculation ^56, 59^ (Figure 5L). These changes were further associated with evidence of ventricular wall defects and altered expression of cell junctional components required for structural integrity of the myocardium ^6, 7, 55^. The disrupted actin cytoarchitecture in early Nrg1 mutants causes a failure to acquire the motile and invasive behaviors required for proper orientation of the mitotic spindle and orthogonal cardiomyocyte division. Later trabecular growth and elongation phases are driven by EMT-like migration of cardiomyocytes derived from the growing outer compact layer towards the trabeculae (Figure 5L).

The initial thickening of the inner myocardial wall therefore depends on the formation of a mixed trabecular/compact zone closest to the source of Nrg1. The outer compact myocardium initially consists of just 2-3 layers, but attains a thickness of about 15-20 cell layers by the end of trabeculation (Figure 5L), likely through differential growth ^96^. We surmise that the inner compact myocardium acts as a “feeder” layer for trabecular growth, ^6^consistent with previous analyses identifying clones of migratory cells spanning anatomical transmural locations during ventricular chamber formation ^6, 8, 95, 97, 98^. These EMT-like processes involve disruption of cell–cell interactions and disassembly of the polarity complexes, with rearrangement of actin filaments to bestow invasive behaviors required for cellular motility ^99^. However, they take place in the absence of the activation of canonical EMT transcription factors or the acquisition of mesenchymal-specific markers and do not involve the establishment of a front-to-back polarity, suggesting that this type of EMT is partial or incomplete ^99^. Thus, Nrg1 regulates the spatial distribution of compact and trabecular myocardium by generating molecularly distinct myocardial layers and establishing the boundaries of trabecular identity (Figure 5L).

Consistent with this notion, by promoting the development of the trabecular and the adjacent inner compact myocardium, Nrg1 restricts the growth of the proliferative cardiomyocytes of the outer layer. By analogy with the situation in lung airway epithelium cell division ^100^, parallel cardiomyocyte division may be the default orientation for myocardial growth. Nrg1-ErbB2/4-Erk1/2 signaling activity may function as a switch to override this default orientation, although not by providing any orientation cue, but rather by determining whether or not a cell responds to such cues. These could be provided by cell polarity pathways, as they play important roles in regulating mitotic spindle angle distribution in epithelial tissues ^60^. We can surmise that, in addition to transcriptional regulatory mechanisms, a Nrg1-pERK kinase cascade may exert effects (direct and indirect) on the centrosomal components of the mitotic spindle, or the cytoskeletal elements that control centrosome position and spindle orientation ^101^.

Nrg1 overexpression causes a profound change in compact ventricular wall composition, increasing the proportion of endothelial and epicardial cells relative to that of cardiomyocytes. This suggests that, as development advances, patterning and growth of the outer compact layer no longer depends on Nrg1 ^102^, as cardiomyocytes are no longer sensitive to signals emanating from the endocardium, and the activation of the downstream effector pErk is suppressed in the myocardium, likely through downregulation of the ErbB2/4 receptors. This would be consistent with a spatio-temporal range of action confined to the trabecular period of ventricular wall development. Consequently, by virtue of their comparatively higher proliferation rate, compact layer cardiomyocytes eventually outgrow trabecular cardiomyocytes during the compaction phase ^13^. Specification to trabecular versus compact myocardium may therefore depend on the relative distance of cardiomyocytes from the source of Nrg1, as its influence becomes limited to the endocardium-proximal cardiomyocytes giving rise to the subendocardial ventricular myocardium and Purkinje fiber conduction system network ^79^ (Figure 5L).

Our studies underscore the tight coordination between cell division and migration that evolved to ensure pattern and boundary formation during tissue growth and morphogenesis ^103–105^. In addition to the morphogenic EMT-like component, Nrg1 regulates gene signatures for cell-cycle progression linked to mitosis, DNA replication and repair, and the cytoskeleton ^106–108^. There is a close association between cytoskeleton organization, cell adhesion, and the cell cycle ^109^, revealed by actin-regulated G2/M checkpoint controls^110–112^, and our interpretation of the mutant phenotype is plausible if Nrg1-pErk signaling is required for progression through the G2/M phase. Consistent with this idea, ERBB2 (HER2) and ERK1/2 have been identified as important regulators of the G2/M checkpoint response to genotoxic stress ^113, 114^. Further, these studies highlight 2 types of cell cycle arrest ^84, 115^: 1) classical (hypomitogenic) growth arrest caused by growth factor withdrawal, is an exit from the cell cycle characterized by the inactivation of both upstream (Ras-Mapk signaling) and downstream proliferation factors (cyclins); 2) hypermitogenic cell cycle arrest caused when mitogen-activate pathways are active but cell cycle progression is blocked downstream (for example p21/p27 over-expression), ultimately leading to a senescence-like phenotype.

With regard to Nrg1-pErk and Hippo-Yap1 interactions, available evidence indicates that: 1) Nrg1 activates endogenous ErbB4 to induce Yap1 target genes in breast cancer cells ^75^; 2) ErbB2 drives mitosis during cardiac regeneration via pErk-dependent and Hippo-independent Yap1 activation ^89^; 3) Interaction of Yap1 with the Myb-MuvB (MMB) complex promotes mitotic gene expression in fetal cardiomyocytes ^90^; and 4) Yap1-mediated cardiac regeneration requires the transcription of cell-cycle, adherens-junction, and cytoskeletal genes ^92, 116–118^. Considering this, Yap1 could act as a downstream effector of Nrg1 during trabeculation (Figure 5L) as there exists a significant overlap between Nrg1, ErbB2 and YAp1 GOF, and Lin9 inactivation gene signatures, involving EMT-like genes, and we provide evidence suggesting that Nrg1 is required for Yap1(S274) nuclear localization during early trabeculation but not during compaction, consistent with a restricted Nrg1-pErk window of function. Intriguingly, the Yap1 homolog WWTR1 promotes compact wall development by inhibiting trabeculation in zebrafish, whereas Nrg1-ErbB2 signaling negatively regulates WWTR1 nuclear localization and activity ^31^. These findings provide incentives for further studies into the role of Nrg1-Yap1 in ventricular wall development.

Our studies provide evidence that non-compaction/hypertrabeculation is a defect of asymmetric ventricular wall growth. Because of the considerable overlap between trabeculation and ventricular wall maturation and compaction, it has been difficult to distinguish excessive trabeculation from the failure to compact and to determine whether these phenotypes form part of a continuum or are separate entities. Experimental evidence from human induced pluripotent cell models suggests that non-compaction might be attributable to defective cardiomyocyte proliferation ^119^, whereas data from mice show that the persistence of trabeculae postnatally may be the result either of excessive proliferation of trabecular cardiomyocytes ^120, 121^ or of reduced proliferation in compact myocardium ^13, 20, 122^. Our studies help to resolve this conundrum by suggesting that non-compaction/hypertrabeculation is linked to disrupted asymmetric growth and differentiation across the ventricular wall, resulting in defective trabecular patterning and cardiomyocyte proliferation. The extensive overlap between cell division and morphogenesis makes it difficult to discern the individual contributions of these processes to trabecular and compact myocardium growth. The possibility of multi-omic measurement at single-cell scale provides an incentive for future investigation.

## Supporting information

Supplemental Material_Grego-Bessa et al

## Nonstandard Abbreviations and Acronyms

4-OHT: 4-hydroxytamoxifen
A-B polarity: apico basal polarity
CM: compact myocardium
EMT-like: epithelial-mesenchymal transition-like
ErbB2/4: erb-b2 receptor tyrosine kinase 2/4
Erk: extracellular signal regulated MAP kinase
GOF: gain-of-function
GSEA: gene set enrichment analysis
IF: immunofluorescence
LOF: loss-of-function
Nrg1: neuregulin-1
OCD: oriented cell division
OE: overexpression
SMA: smooth muscle actin
SM22: smooth muscle cell-specific cytoskeletal protein
TM: trabecular myocardium
VCS: ventricular conduction system
Yap1: yes1-associated transcriptional regulator

## Acknowledgments

We thank A. Galicia and L. Méndez for mouse husbandry, the Genomics Units for carrying out RNA-seq, and S. Bartlett for English editing.

