## Supplemental Material_Grego-Bessa et al for "Neuregulin-1 regulates cardiomyocyte dynamics, cell cycle progression, and maturation during ventricular chamber morphogenesis"

### ONLINE SUPPLEMENTARY DATA

----------------------------------------------------------------------------------------------------------------

### DETAILED METHODS

#### Mouse strains and genotyping

To study the role of Nrg1 signaling in cardiac development, we used a conditional *Nrg1$^{flox}$* allele [1] and the following CRE driver mouse strains: *Tie2$^{Cre}$* [2], *Pdgfb-iCre$^{ERT2}$* [3], *Mesp1$^{Cre}$* [4], *Cdh5$^{CreERT2}$* [3], and *;Nkx2-5$^{Cre}$* [5]. Mouse lines were genotyped using the primers listed in Table S5. Animal studies were approved by the CNIC Animal Experimentation Ethics Committee and by the Community of Madrid (Ref. PROEX 155.7/20). All animal procedures conformed to EU Directive 2010/63EU and Recommendation 2007/526/EC regarding the protection of animals used for experimental and other scientific purposes, enacted in Spanish law under Real Decreto 1201/2005.

#### Generation of *R26PA-Nrg1-iresGFP* transgenic mice

A full-length cDNA from mouse *Nrg1* (2103 bp) was obtained from clone IMAGE 100064088. The sequence was PCR-amplified with Phusion High-Fidelity DNA Polymerase (NEB). The PCR product was digested with *Nhe*I and cloned into a *pBigT-IRESeGFP* plasmid previously generated by cloning a *Sal*I *IRES-eGFP* fragment into the *Xho*I site of *pBigT*. The resulting *loxP-PGK-Neo*-3X-STOP-*loxP*-*Nrg1-IRES-eGFP* fragment was cloned into the *Pac*I and *Asc*I sites of the *pROSA26PA* plasmid [6] (Figure S13A). Gene targeting of this construct was performed in G4 mouse embryonic stem cells (mESCs) and confirmed by Southern blotting with external 5′ and 3′ hybridization probes (Figure S13A,B). Mice were generated by injecting targeted cells into B6CRL blastocysts to generate chimeras that were then analyzed for germline transmission. The selected animals were backcrossed to the C57BL/6 background. For simplicity, we call the line *R26Nrg1$^{GOF}$;Nkx2-5Cre.*

**4-Hydroxytamoxifen induction**

For inducible CRE lines, we induced nuclear CRE by oral gavage with 200 µl of 4-hydroxytamoxifen (4-OHT) (Sigma H6278; 5 mg/ ml solution prepared by diluting 50 mg 4-OHT in 1 ml 95% ethanol plus 9 ml corn oil). Double heterozygous *Nrg1$^{flox/+}$;Cdh5$^{CreERT2/+}$* females were crossed with homozygous *Nrg1$^{flox/flox}$* males, and pregnant females were induced by oral gavage at E10.5 and E11.5 and dissected at E16.5 or were induced at E12.5 and 13.5 and dissected at E18.5. For the analysis of coronary vessels, double heterozygous *Nrg1$^{flox/+}$;Pdgfb1$^{CreERT2/+}$* females were crossed with homozygous *Nrg1$^{flox/flox}$* males, and pregnant females were induced at E10.5 and E11.5 and dissected at E16.5.

**Tissue processing**

Dissected early embryos (E8.0-E10.5) were sorted according to somite number to ensure comparable developmental stages in all experiments with control and mutant embryos. For immunostaining, E8.5 and E9.5 embryos were fixed in 4% paraformaldehyde (PFA) for 2 hours at 4ºC. E10.5 and older embryos were fixed overnight (O/N) at 4ºC. For studies involving *in situ* hybridization (ISH) or staining with hematoxylin and eosin (H&E) or Alcian blue, all embryos were fixed O/N. After dehydration through a graded ethanol series followed by xylene washes, embryos were embedded in paraffin at 65ºC. For cryosections, embryos were fixed in 4% PFA, washed in 30% sucrose, embedded in optimal cutting temperature (OCT) compound and stored at -80ºC.

**Histology, Alcian blue staining, and *in situ* hybridization**

H&E and Alcian blue stainings were performed on sections according to standard protocols. ISH was performed as described [7, 8]. Details of probes will be provided on request.

**Bright field images**

Images of H&E, Alcian blue, and ISH stainings were acquired with an Olympus BX51 fluorescence microscope coupled to a Nikon DP71 camera and CellSens software.

**Quantification of compact myocardium thickness and trabecular length and width.**

Paraffin sections (7 µm) or cryosections (10 µm) were stained for myocardium- and endocardium-specific markers. Compact myocardium thickness was measured with Fiji software (ImageJ plugin). For E9.5 and E10.5 embryos, trabecula length was measured in 3 non-consecutive sections in 4 ventricular regions of at least 3 *Nrg1$^{flox}$*;*Tie2$^{Cre}$* and 3 control embryos. For E16.5 embryos, trabecula length was measured in 3 non-consecutive sections from 4 ventricular regions from at least 3 embryos per genotype

(*Nrg1*$^{flox/+}$;*Cdh5*$^{CreERT2/+}$, 3 *R26Nrg1*$^{GOF}$;*Nkx2-5*$^{Cre}$, and wild type controls). Mean values are presented in µm.

**Whole-mount immunofluorescence**

Embryos were fixed for 3 hours (E10.5) or O/N (E16.5) at 4ºC in 4% PFA. Fixed embryos were permeabilized by incubation for 1h with 0.5% Triton X-100 in PBS (0.5% PBS-TX) and blocked O/N in 0.5% PBS-TX containing 10% FBS. Embryos were incubated with primary antibodies O/N at 4ºC in 0.5% PBS-TX, 10% FBS. After several washes in 0.5% PBS-TX, 10% FBS, embryos were incubated with secondary antibodies in the same solution O/N at 4ºC. After several washes in PBS containing 0.1% Tween-20 (PBS-T), embryos were mounted on a slide. To image chamber interiors, E10.5 and E16.5 hearts were dissected and placed on a petri dish in PBS, and a piece of the ventricular wall was then cut to visualize the trabecular network. Samples were mounted on a slide between two strips of tape separated by 0.5 cm to create a 3D space, allowing conservation of the original sample shape [9]. Tissue was mounted in Vectashield medium and covered with a cover slide. For procedure details see [10]. The coronary vasculature was visualized as described [11]. Measurements were taken with ImageJ software. The main coronary trees were selected manually with the 'Freehand selection' tool and measured directly from the autoscaled images obtained by Z-projection. The selected area was quantified in mm$^2$.

**Immunofluorescence on sections**

Paraffin-embedded 7-µm sections or OCT-embedded 10-µm sections were blocked for 30 min with 0.3% PBS-TX, 5% FBS and incubated O/N at 4ºC with primary antibodies diluted in the same solution (Table S5). Sections were then washed several times in 0.3% PBS-TX, 5% FBS and incubated for 1h with an appropriate fluorescent-dye-conjugated secondary antibody diluted in the same solution (Table S5). Specific immunostainings were performed using tyramide signal amplified (TSA) coupled to a fluorophore (Akoya Biosciences, Del Monte *et al.*, 2011). BrdU immunostainings were amplified with a biotinylated secondary antibody, ABC (Vectastain Kit), and TSA. Nuclei were counterstained with DAPI.

**Fluorescence signal quantification**

Pixel intensity was measured on *en face* images of whole-mount stained ventricles. Data were obtained from Z-stack projections of 4 optical sections taken every 1 µm with a Leica SP8 confocal microscope. Green signal was transformed to gray scale with Fiji. Pixel intensity was plotted along 5-pixel wide 50 mm lengths. Data are expressed as gray scale intensity (0 to 255) .

**Proliferation analysis and quantification**

Cell proliferation was evaluated as the incorporation of 5-bromo-2'-deoxyuridine (BrdU) into DNA and the immunodetection of phospho-histone H3 immunodetection. Pregnant females received intraperitoneal injections of 200 µl BrdU (10 mg/ml). After 2h, embryos were collected in PBS and processed for immunofluorescence detection with a rat anti-BrdU monoclonal antibody (Table S5, sheet 2) followed by a anti-rat-biotinylated secondary antibody (Table S5, sheet 3). The signal was amplified with TSA technology (Akoya Biosciences). The total numbers of DAPI- and BrdU-positive cells in the myocardium and endocardium of developing hearts were counted on non-consecutive sections (≥ 3) in at least 3 embryos using Fiji software (Image J plugin).

**Quantification of cell roundness**

Cryosections stained with wheat germ agglutinin (WGA) coupled with FITC or Rhodamine (Table S5, sheet 2) were imaged with a Leica SP8 confocal microscope fitted with a 63x lens. Cells were delineated with the Freehand Selection Tool and analyzed with the Shape Descriptors Tool in Fiji/Image J plugin. Several hundred cells were analyzed from 3 biological replicates. Results are expressed in $\mu m^2$.

**Quantification of oriented cell division**

Yellow lines were drawn to indicate the orientation of division for each cardiomyocyte, and white lines were drawn in parallel to the basement membrane and cardiac lumen for each analyzed cardiomyocyte. The plane of cell division was measured as the angle between the yellow and white lines. Division planes forming angles between 70º and 90º to the basement membrane were classified as perpendicular, those between 20º and 70º as oblique, and those between 0º and 20º as parallel. Measurements were made with the Fiji ImageJ plugin. Mitotic cardiomyocytes (MC, α-SMA-positive) were analyzed in the epithelial myocardium at E8.5 or in compact myocardium at E9.5. Cardiomyocytes of at least 3 embryos were analyzed at cell-cycle phases when the orientation of the division is easily identifiable (anaphase, telophase, or cytokinesis) and was also detected by survivin immunofluorescence.. Results are shown as the percentage of each type of division (parallel, oblique, and perpendicular) at E8.5 and E9.5. *P* values were obtained by Fisher's exact test analysis from the total MC analyzed.

**Isolectin B4 quantification**

The area positive for integrin Isolectin B4 (IB4) immunofluorescence staining was measured with the Fiji/Image J plugin. The myocardial region of interest was selected as the area positive for the myocardial marker α-SMA, and a threshold was established to select the area positive for the IB4 signal (grey signal within the myocardial region). This was done in 3 embryos per genotype, in 3 non-consecutive sections, and in 3 myocardial regions per section (from the right, medium, and left ventricular regions). Results are expressed in % of IB4+ signal in compact myocardium.

**Integrin α6 quantification**

The area positive for integrin α6 (ITGα6) immunofluorescence staining was measured with the Fiji/Image J plugin. The myocardial region of interest was selected as the area positive for the myocardial marker α-SMA, and a threshold was established to select the area positive for the ITGα6 signal (green signal within the myocardial region). This was done in 3 embryos per genotype, in 3 non-consecutive sections, and in 3 myocardial regions per section (from the right, medium, and left ventricular regions). Results are expressed in $\mu m^2$.

**N-cadherin quantification**

The WGA signal was used as a reference of the cardiomyocyte membrane along the basal region on each section. Cardiomyocytes with N-cadherin localized to the basal region, facing the cardiac lumen, were counted in 3 control and 4 $Nrg1^{flox};Tie2^{Cre}$ embryos, in 3 non-consecutive sections per embryo and in 3 compact myocardium regions per section (from the right, medium, and left ventricular regions). Results are presented as the percentage of cardiomyocytes with basal N-cadherin expression.

**Confocal imaging and 3D reconstruction**

Confocal images of whole embryos and tissue sections were acquired with a Nikon A1R laser scanning confocal microscope fitted with a 10X, 20X, or 60X objective; a Leica SP8 confocal microscope fitted with a 20X or 63X objective; and a Zeiss 780 confocal microscope fitted with a 20X objective with a dipping lens. Images were viewed and processed with NIS-Elements SD Imaging Software. Images of whole-mount embryos were collected as Z-stacks of optical sections taken every 2 µm (E10.5 hearts) or every 1-10 µm (E16.5 hearts). Z-projections and 3D images were assembled using Fiji/ImageJ plugin and IMARIS software (Bitplane Scientific Software). Images were processed in Adobe Photoshop Creative Suit 5.1 and Fiji/ImageJ plugin.

**Western blotting**

Protein extracts were obtained from ventricles of E9.5 or E16.5 embryonic hearts in 20 µl or 100 µl of Tissue Protein Extraction Reagent (T-PER, Thermo Fisher Scientific #78510) supplemented with Halt Phosphatase Inhibitor (100X; Thermo Scientific #74827) plus Complete Protease Inhibitor Cocktail (Roche #11697498001, Germany). Western blots were performed according to standard protocols, and proteins were detected with HRP-conjugated secondary antibodies and ECL detection reagents (Amersham, UK).

**RNA-Seq**

Whole hearts were harvested from E9.5 $Nrg1^{flox};Tie2^{Cre}$ mice. Ventricles were isolated from E15.5 $Nrg1^{flox};Cdh5Cre^{ERT2}$ and $R26Nrg1^{GOF};Nkx2\text{-}5^{Cre}$ mice. For whole hearts (n=4 replicas, consisting of 6 pooled hearts for each control and mutant), RNA was extracted using Arcturus PicoPure RNA isolation kit (Thermo Fisher Scientific). For ventricles (n=4 replicas, consisting of 1 ventricle for each control and mutant/transgenic), RNA was extracted using RNeasy mini plus kit (Qiagen). Total RNA (100-200ng) was used to generate barcoded RNA-seq libraries using the NEBNext Ultra RNA Library preparation kit (New England Biolabs). Briefly, poly A+ RNA was purified using poly-T oligo-attached magnetic beads followed by fragmentation and then first and second cDNA strand synthesis. Next, cDNA 3' ends were adenylated and the adapters were ligated followed by PCR library amplification. Finally, the size of the libraries was checked using the Agilent 2100 Bioanalyzer DNA 1000 chip and their concentration was determined using the Qubit® fluorometer (Life Technologies). Libraries were sequenced on a HiSeq2500 or HiSeq4000 (Illumina) to generate 60 bases single reads. FastQ files containing reads for each library were extracted and demultiplexed using bcltofastQ (Illumina). Sequencing adapter contaminations were removed with cutadapt, discarding those reads that were shorter than 30 nt after trimming. Resulting reads were aligned against mouse reference transcriptome GRCm38, Ensembl genebuild 91, and gene expression was quantified using RSEM v1.2.3 [12]. Differential gene expression was tested using a generalized linear model as implemented in the EdgeR package from Bioconductor [13]. Counts were normalized by the TMM method. Genes with at least 1 cpm in at least 3 samples were defined as expressed and were retained for later analysis. Genes showing altered expression with an FDR < 0.05 were considered differentially expressed. ClustVis [14] and R were used to generate heatmaps for selected collections of differentially expressed genes. Functional analyses were performed with GSEA [15] on the complete set of expressed genes against Hallmark gene sets, and Panther [16] on the collections of differentially expressed genes.

Data are deposited in the NCBI GEO database under accession number GSE216471. The following secure token has been created to allow reviewer access while it remains in private status: ihsvsqyobpijnev

**Statistics**

Statistical comparisons were made by unpaired two-tailed Student *t*-test in Graph Pad (Prism 5.0). Data are presented as mean ± S.D. unless otherwise indicated. Differences were considered statistically significant at P <0.05. Categorical comparisons were made by Fisher's exact test in Graph Pad (Prism 5.0). All statistics are shown in Table S6.

**Supplementary Figure Legends**

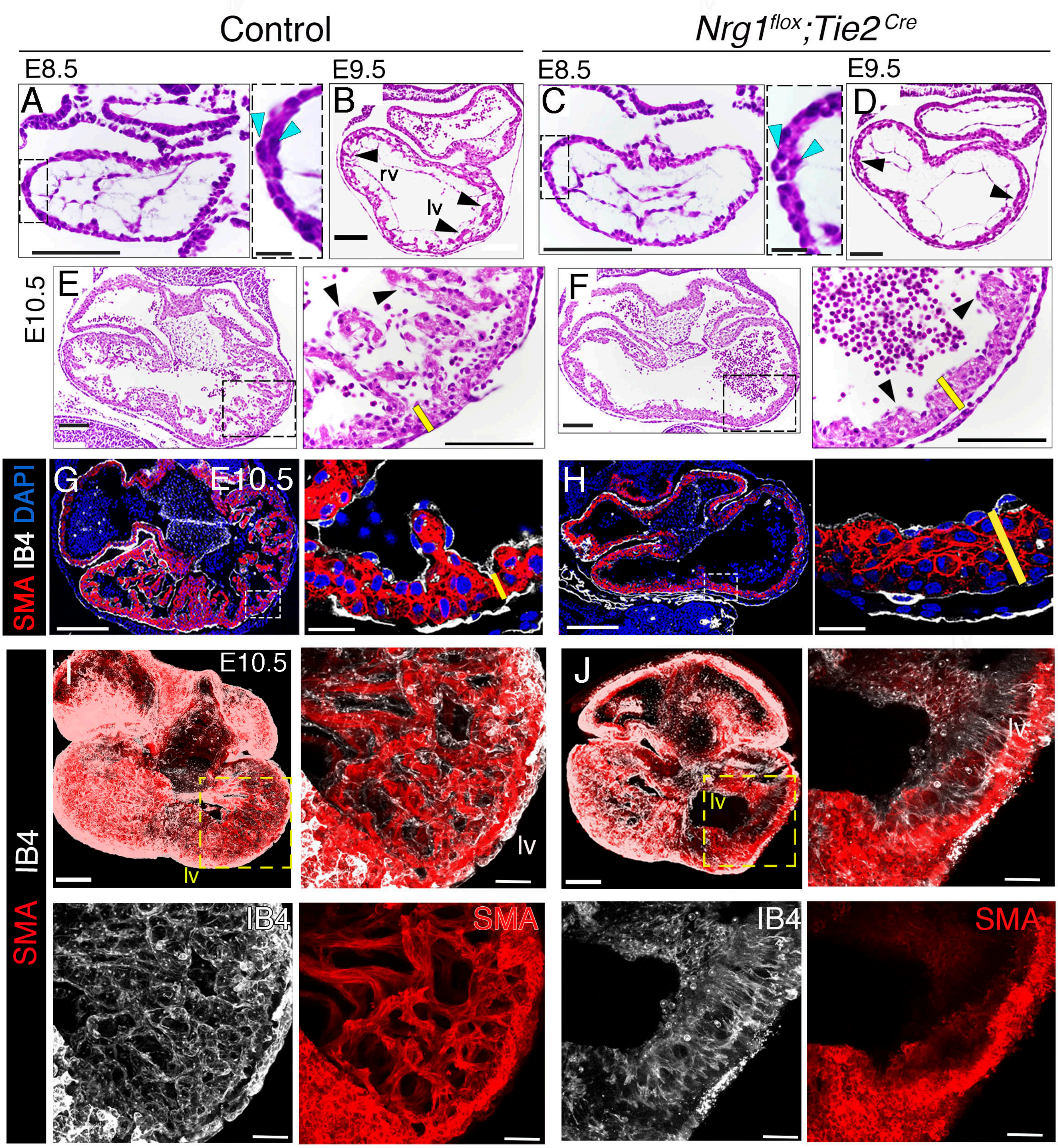


Figure S1_Grego Bessa

**Figure S1. Histological analysis reveals impaired trabeculation in *Nrg1$^{flox}$;Tie2$^{Cre}$* mutant hearts. (A-F)** H&E staining of E8.5, E9.5 and E10.5 control **(A, B, E)** and *Nrg1$^{flox}$;Tie2$^{Cre}$* heart sections **(C, D, F)**. Blue arrowheads in insets mark nuclei in compact myocardium; black arrowheads mark trabeculae. Yellow bars indicate compact myocardium thickness **(G, H)** Immunofluorescence against smooth muscle actin (SMA, red) and isolectin B4 (IB4, white) in heart sections from E9.5 control **(G)** and *Nrg1$^{flox}$;Tie2$^{Cre}$* embryos **(H)**. Yellow lines mark the thickness of the compact myocardium (CM). Sections were counterstained with DAPI (blue). **(I, J)** 3D reconstruction of 50-µm thick sections of E10.5 control and *Nrg1$^{flox}$;Tie2$^{Cre}$* heart sections stained for smooth muscle actin (SMA) and isolectin B4 (IB4). lv, left ventricle. Scale bars, 100 µm in A-F general views and details of ventricles; 200 µm in G-J; 100 µm in I,J magnifications; 20 µm in G,H magnifications.

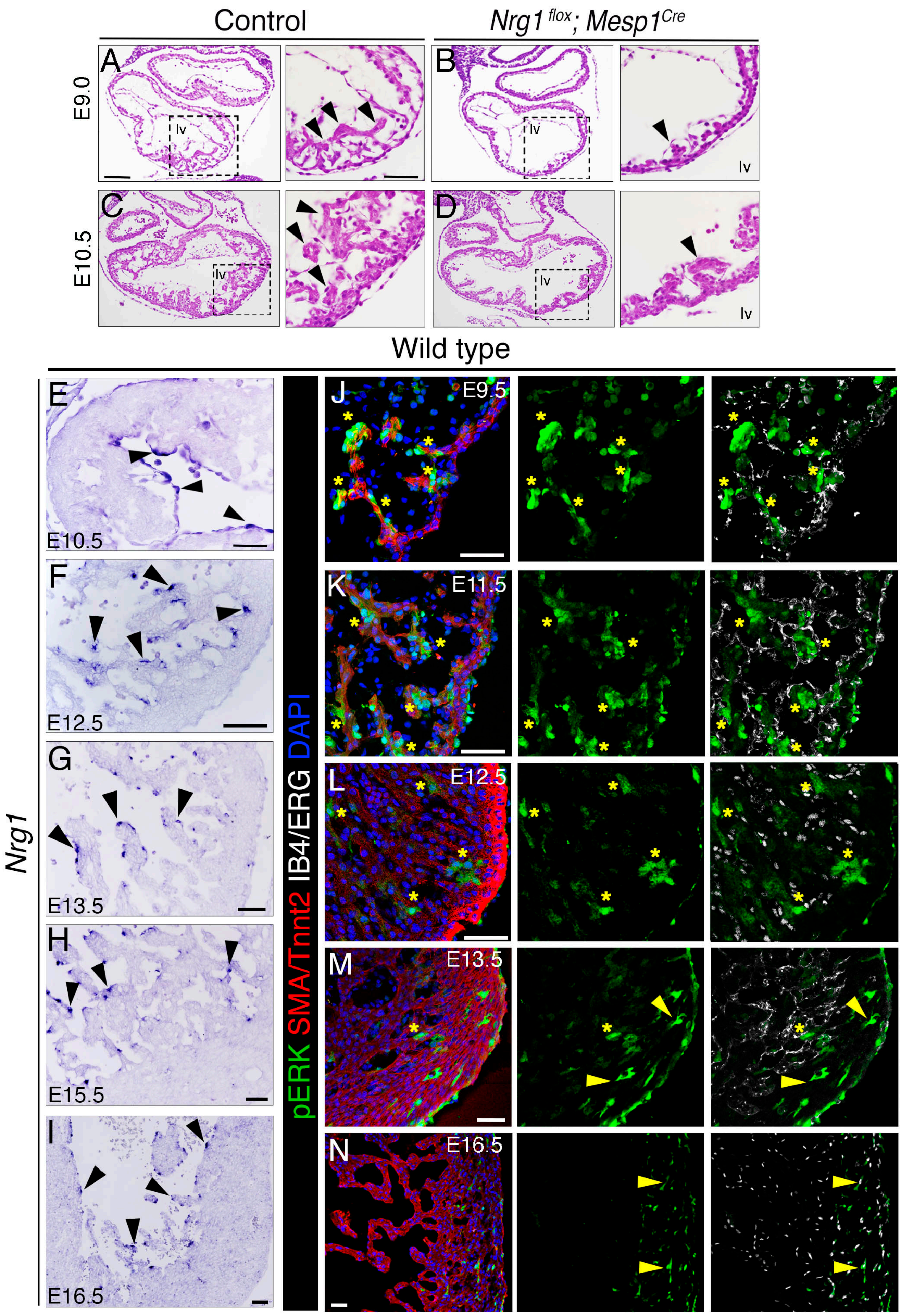


Figure S2_Grego-Bessa

**Figure S2. Defective trabeculation in *Nrg1$^{flox}$;Mesp1$^{Cre}$* mutants. pERK expression in the developing ventricle. (A-D)** H&E staining of E9.5 and E10.5 control and *Nrg1$^{flox}$;Mesp1$^{Cre}$* heart sections. Arrowheads mark trabeculae. **(E-I)** *Nrg1* ISH in E10.5, E12.5, E13.5, E15.5 and E16.5 wild type hearts. The arrowheads point to *Nrg1*-expressing endocardial cells. (**J-N**) pERK staining in transverse sections of E9.5, E11.5, E12.5, E13.5, and E16.5 wild type hearts. Asterisks mark pERK-positive cardiomyocytes; arrowheads mark pERK-positive endothelial cells. The myocardium was stained by immunofluorescence against SMA (J-M) or Tnnt2 **(N)**. Endocardium was immunostained for isolectin B4 (IB4, J,K,M) or ERG (L,N). Scale bars, 100 μm in A-D general views, 50 μm in magnifications; 50 μm in E-N.

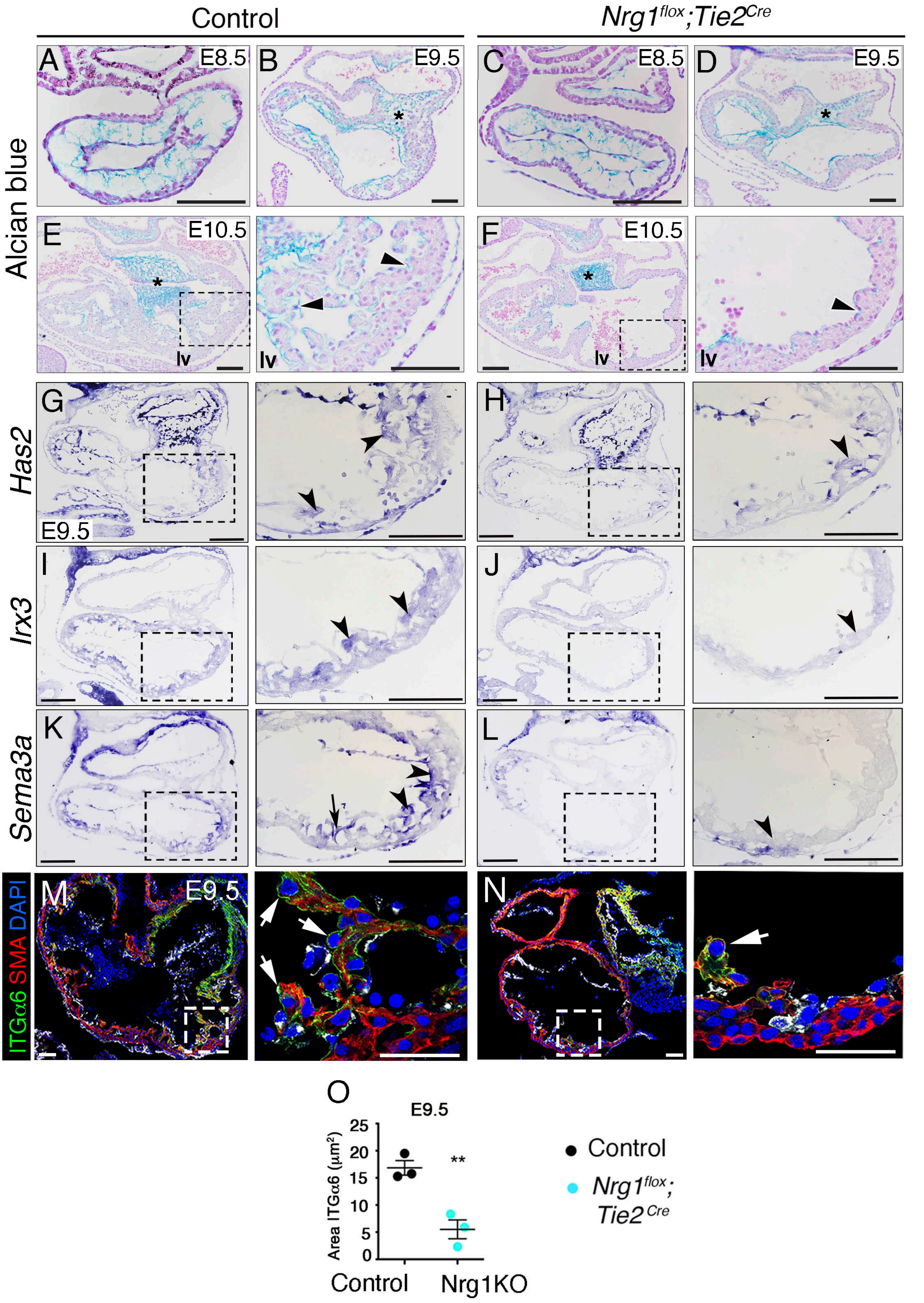


Figure S3_Grego-Bessa

**Figure S3. Reduced extracellular matrix deposition in *Nrg1$^{flox}$;Tie2$^{Cre}$* mutant hearts. (A-F)** Alcian blue staining of glycosaminoglycan on heart sections from E9.5, E10.5, and E11.5 control **(A, B, E)** and *Nrg1$^{flox}$;Tie2$^{Cre}$* embryos **(C, D, F)**. Asterisks mark the atrioventricular canal, arrowheads the trabecular extracellular matrix (ECM). **(G-L)** ISH analysis of *Has2, Irx3*, and *Sema3a* on heart sections from control **(G,I,K)** and *Nrg1$^{flox}$;Tie2$^{Cre}$* embryos **(H,J,L)**. Arrowheads mark trabeculae, arrows endocardium. **(M,N)** Immunofluorescence against integrin α6 (Itgα6) in E9.5 control and *Nrg1$^{flox}$;Tie2$^{Cre}$* heart sections. SMA (smooth muscle actin) **(O)** Quantification of the Itgα6 immunofluorescence area (n=3). **$P$ < 0.01; Student t-test. Scale bars, 100 µm in A-L; 50 µm in M,N.

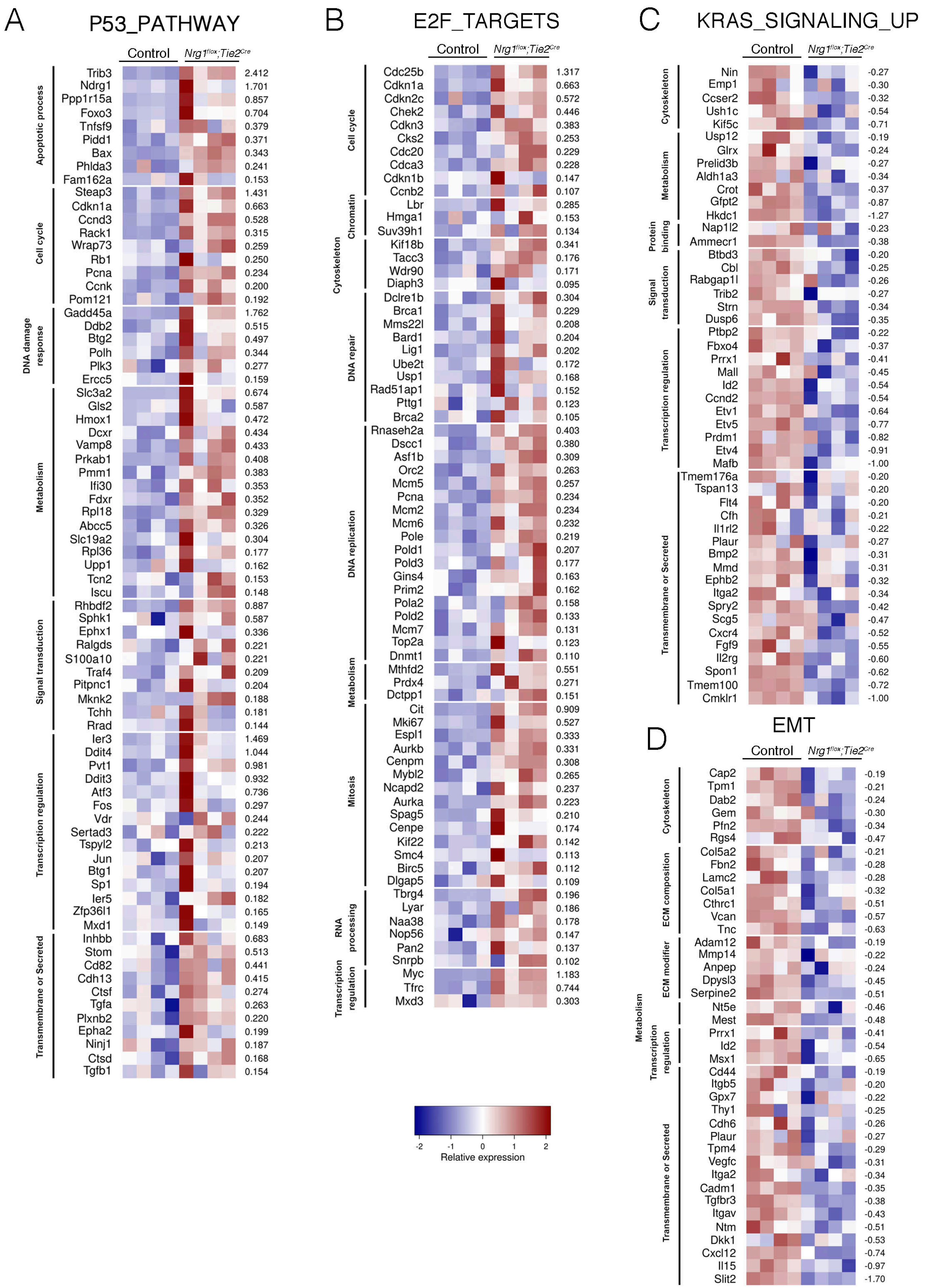


Figure S4_ Grego-Bessa

**Figure S4. GSEA analysis (HALLMARK gene sets) of E9.5 Control vs. *Nrg1$^{flox}$;Tie2$^{Cre}$* heart expression profiles.** Heatmaps represent relative expression values, in each sample, for the "leading edge" genes associated with gene sets (**A**) P53_PATHWAY, (**B**) E2F_TARGETS, (**C**) KRAS_SIGNALING_UP, and (**D**) Epithelial to mesenchyme transition (EMT), extracted from results described in Figure 1. Genes were manually classified into functional categories according to their description in GeneCards (https://www.genecards.org/). Numbers on the right represent logFC values.

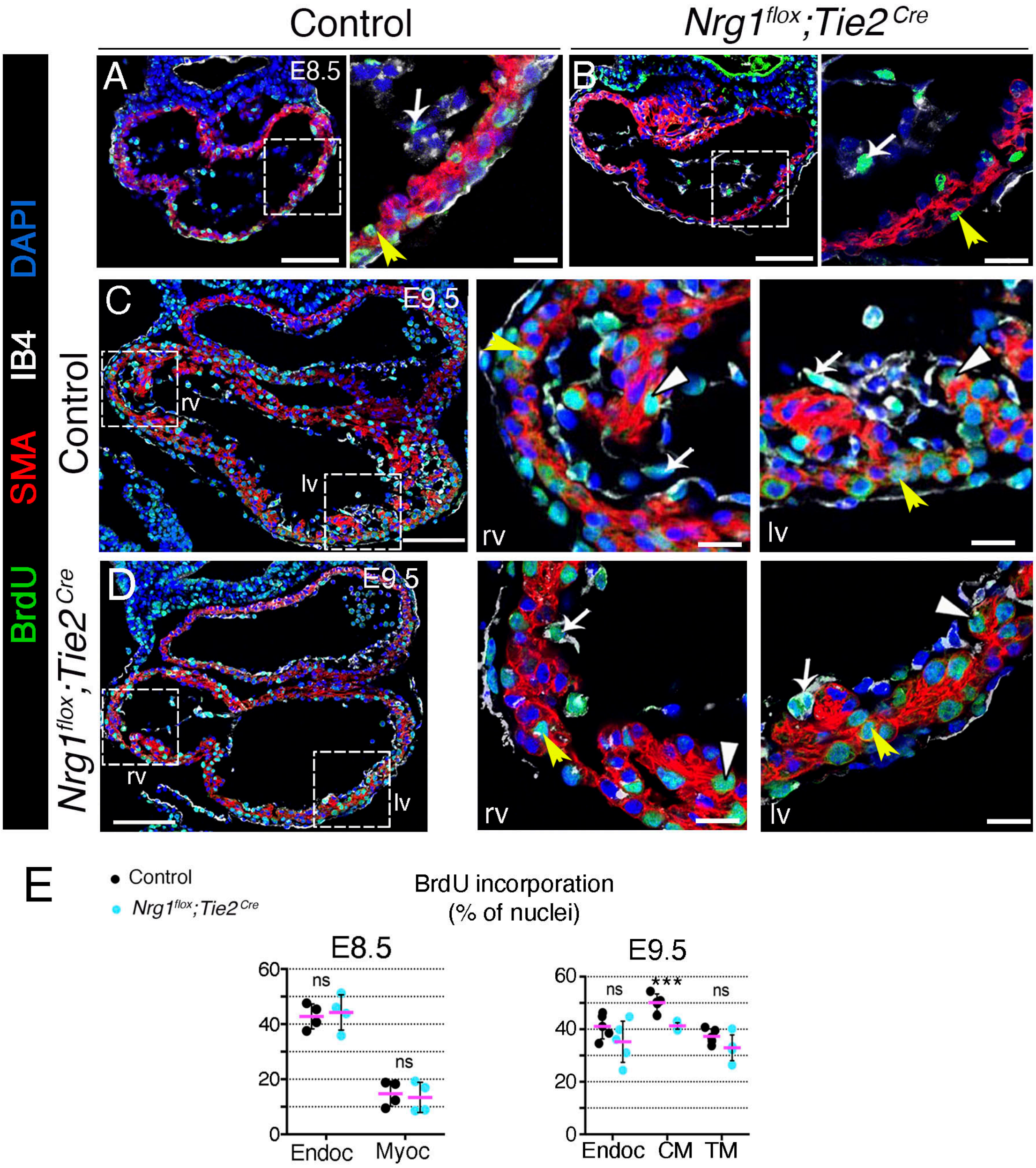


Figure S5_Grego-Bessa

**Figure S5. Reduced cardiomyocyte proliferation in *Nrg1$^{flox}$;Tie2$^{Cre}$* mutants. (A-D)** BrdU immunodetection on heart sections from E8.5 and E9.5 control **(A, C)** and *Nrg1$^{flox}$;Tie2$^{Cre}$* embryos **(B, D)**. White arrows mark nuclear staining in endocardium; yellow arrowheads mark nuclear staining in myocardium. **(E)** Quantification of BrdU incorporation (n = 4). *** $P$< 0.001; ns. non-significant; Student t-test. rv (right ventricle); lv (left ventricle). Endoc (endocardium); Myoc (myocardium); CM, compact myocardium; TM, trabecular myocardium. Scale bars, 100 μm in A-D; 25 μm in insets.

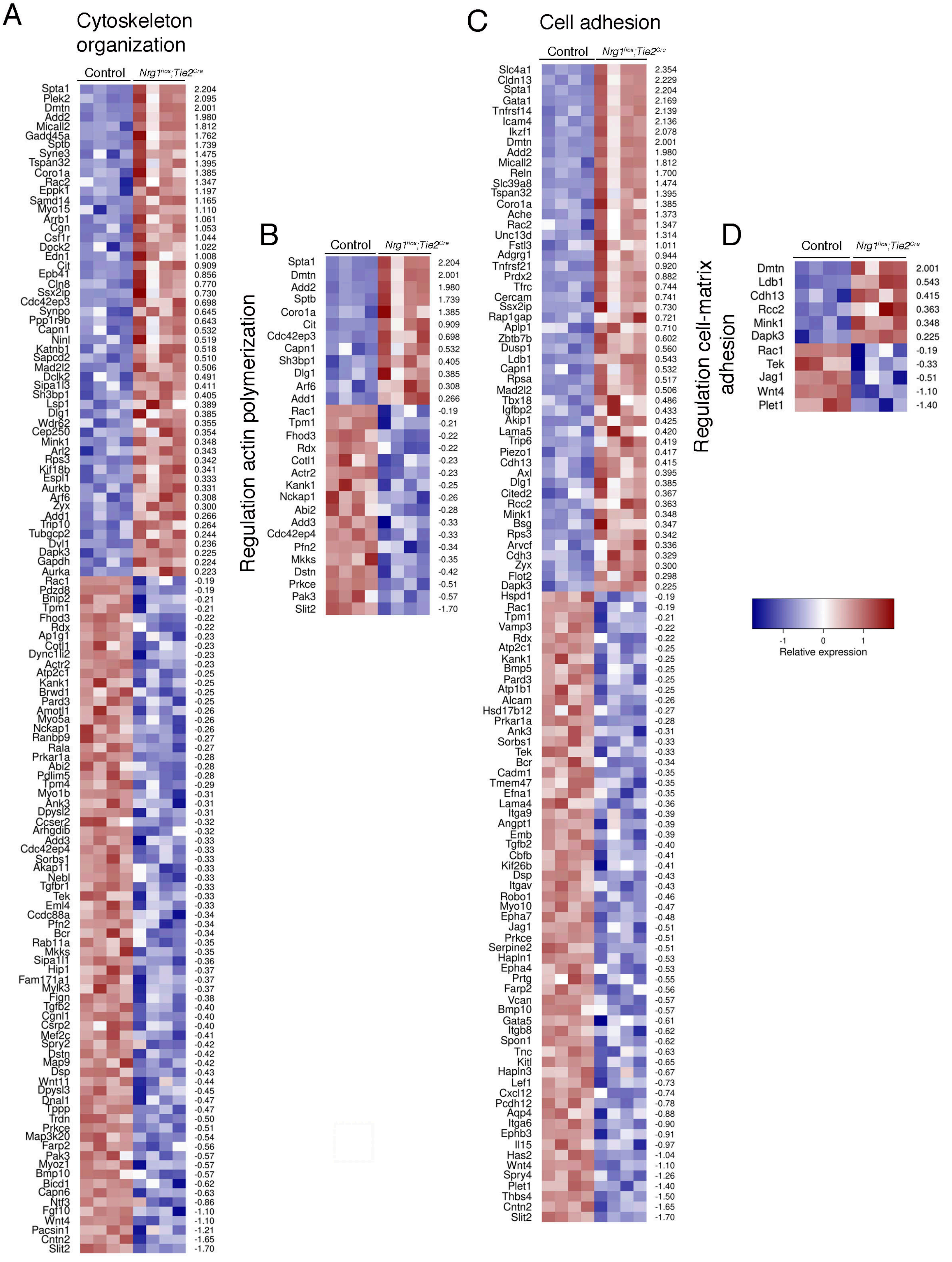


Figure S6_ Grego-Bessa

**Figure S6. PANTHER analysis for E9.5 Control vs. *Nrg1^flox^;Tie2^Cre^*.** The following GO terms were significantly associated to the collection of genes detected as differentially expressed in the E9.5 *$Nrg1^{flox}$;$Tie2^{Cre}$* vs. Control contrast: (**A**) "Cytoskeleton organization" (GO:0007010; pval = 2.98E-10), (**B**) "Regulation of actin polymerization or depolymerization" (GO:0008064; pval = 5.74E-07), (**C**) "Cell adhesion" (GO:0007155; pval = 2.49E-05) , and (**D**) "Regulation of cell-matrix adhesion" (GO:0001952; pval = 3.03E-02). Heatmaps represent relative expression values for differentially expressed genes annotated with each GO term.

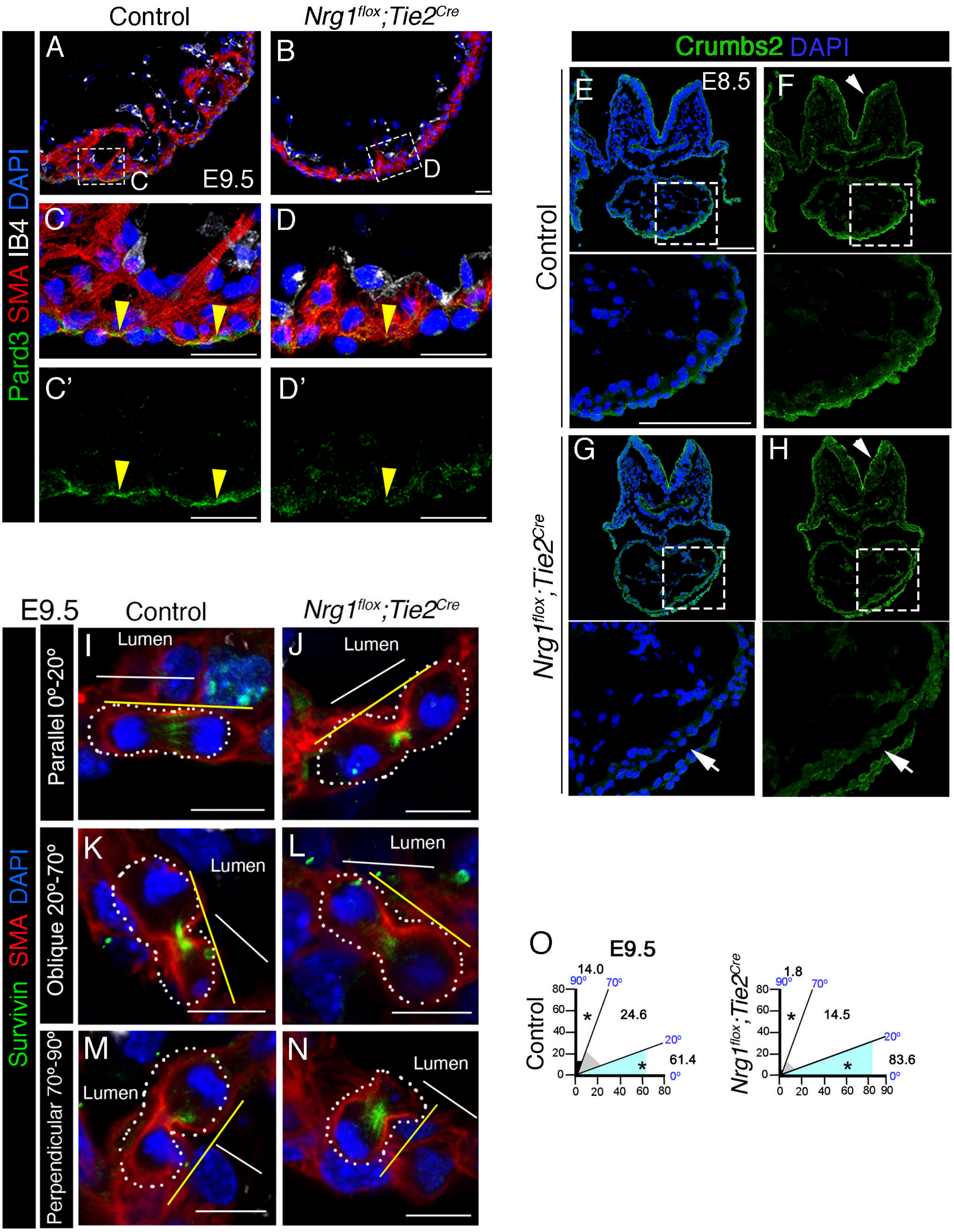


Figure S7_Grego-Bessa

**Figure S7. Disrupted apico-basal polarity in the ventricles of *Nrg1^flox^;Tie2^Cre^* mutants.** Pard3 immunofluorescence on heart sections from E9.5 control embryos **(A,C,C′)** and *Nrg1^flox^;Tie2^Cre^* embryos **(B,D,D′)**. IB4, isolectin B4; SMA, smooth muscle actin. Yellow arrowheads mark Pard3 staining. **(F,G)** Immunofluoresence against Crumbs2 in transverse sections of E8.5 control and *Nrg1^flox^;Tie2^Cre^* embryos (green). DAPI counterstaining is shown in blue. Arrowheads mark the apical domain of the neural tube and indicate similar signal intensities. Arrows mark the apical domain of the compact myocardium. **(I-N)** Representative immunofluorescence images of mitotic cells stained against Survivin (green) and SMA (red) in E9.5 control and *Nrg1^flox^;Tie2^Cre^* heart sections. DAPI counterstaining is shown in blue. White lines indicate the reference plane of the cardiac lumen. Yellow lines indicate the orientation of the mitotic spindle. **(O)** Quantification of OCD in E9.5 control (N=3 embryos; n=57 mitotic figures) and *Nrg1^flox^;Tie2^Cre^* hearts (N=3 embryos; n=45 mitotic figures). *P* values were obtained by Fisher's exact test. **P*-value < 0.05. Scale bars, 25 µm in A-D'; 200 µm in E-H; 10 µm in I-N.

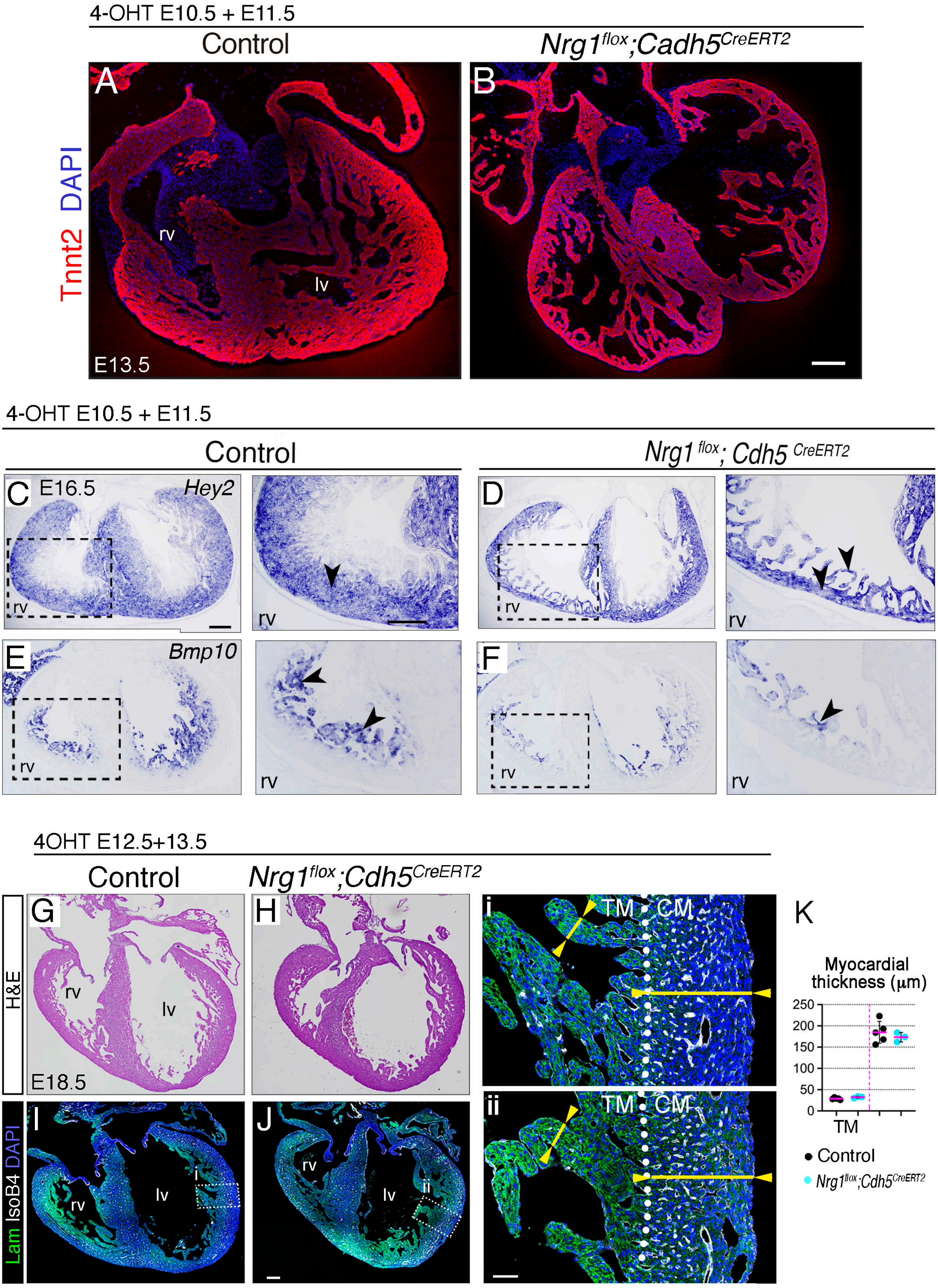


Figure S8_ Grego-Bessa

**Figure S8. Temporal window of Nrg1 requirement and marker analysis in *Nrg1$^{flox}$;Cdh5$^{CreERT2}$* hearts.** **(A)** Troponin T2 (Tnnt2) immunodetection on sections of E13.5 control and *Nrg1$^{flox}$;Cdh5$^{CreERT2}$* hearts **(B)**, after 4-OHT induction at E10.5 and E11.5. Nuclei are counterstained with DAPI. **(C-F)** ISH analysis of *Hey2* and *Bmp10* on E16.5 control and mutant heart sections. The arrowheads mark intense staining. **(G-J)** E18.5 hearts after 4-OHT induction at E12.5 and E13.5. H&E staining (**G,H**) or laminin (Lam) and IsoB4 immunostaining (**I,J**). (**i,ii**) insets: The white dotted lines denote the anatomical border between compact myocardium (CM) and trabecular myocardium (TM). The yellow line highlights CM and TM thickness. rv (right ventricle); lv (left ventricle). **(K)** Quantification of CM and TM thickness in E18.5 hearts after 4-OHT induction at E12.5 and E13.5 ($n \geq 3$). rv, right ventricle; lv, left ventricle. Scale bars, 100 μm in A-F; 200 μm in G-J; 50 μm in i,ii.

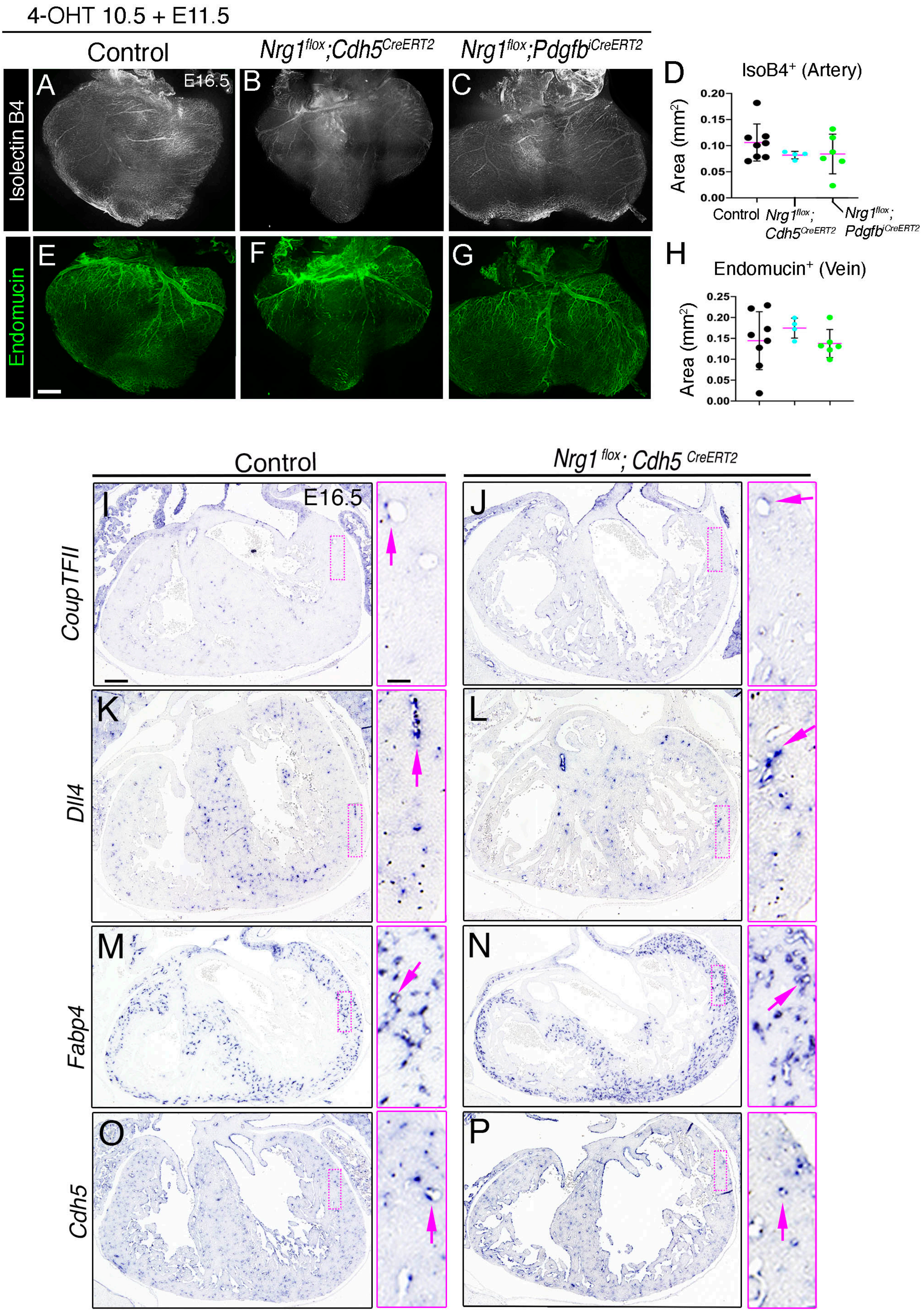


Figure S9_Grego-Bessa

**Figure S9. Coronary vessel development does not depend on Nrg1 signalling. (A-C)** Dorsal views of immunofluorescence staining for Isolectin B4 in whole-mount hearts from E16.5 control **(A)**, *Nrg1*$^{flox}$*;Cdh5*$^{CreERT}$ **(B)**, and *Nrg1*$^{flox}$*;Pdgfb-iCre*$^{ERT2}$ embryos **(C)** after 4-OHT induction at E10.5 and E11.5. **(D)** Quantification of area occupied by IsoB4 (coronary arteries) (n ≥ 3). **(E-G)** Dorsal views of immunofluorescence staining for Endomucin (coronary veins) from E16.5 control **(E)**, *Nrg1*$^{flox}$*;Cdh5*$^{CreERT}$ **(F)**, and *Nrg1*$^{flox}$*;Pdgfb-iCre*$^{ERT2}$ embryos **(G)** after 4-OHT induction at E10.5 and E11.5. **(H)** Quantification of area occupied by endomucin (coronary veins) (n ≥ 3). ns. non-significant; Student t-test. **(I)** ISH analysis of *CoupTFII*, *Dll4*, *Fapb4*, and *Cdh5* on E16.5 control and *Nrg1*$^{flox}$*;Cdh5*$^{CreERT}$ heart sections (4-OHT induction at E10.5 and E11.5). Arrows n insets indicate staining in coronaries. Scale bars, 200 μm in A-G; 100 μm in I-P; 22 μm in insets.

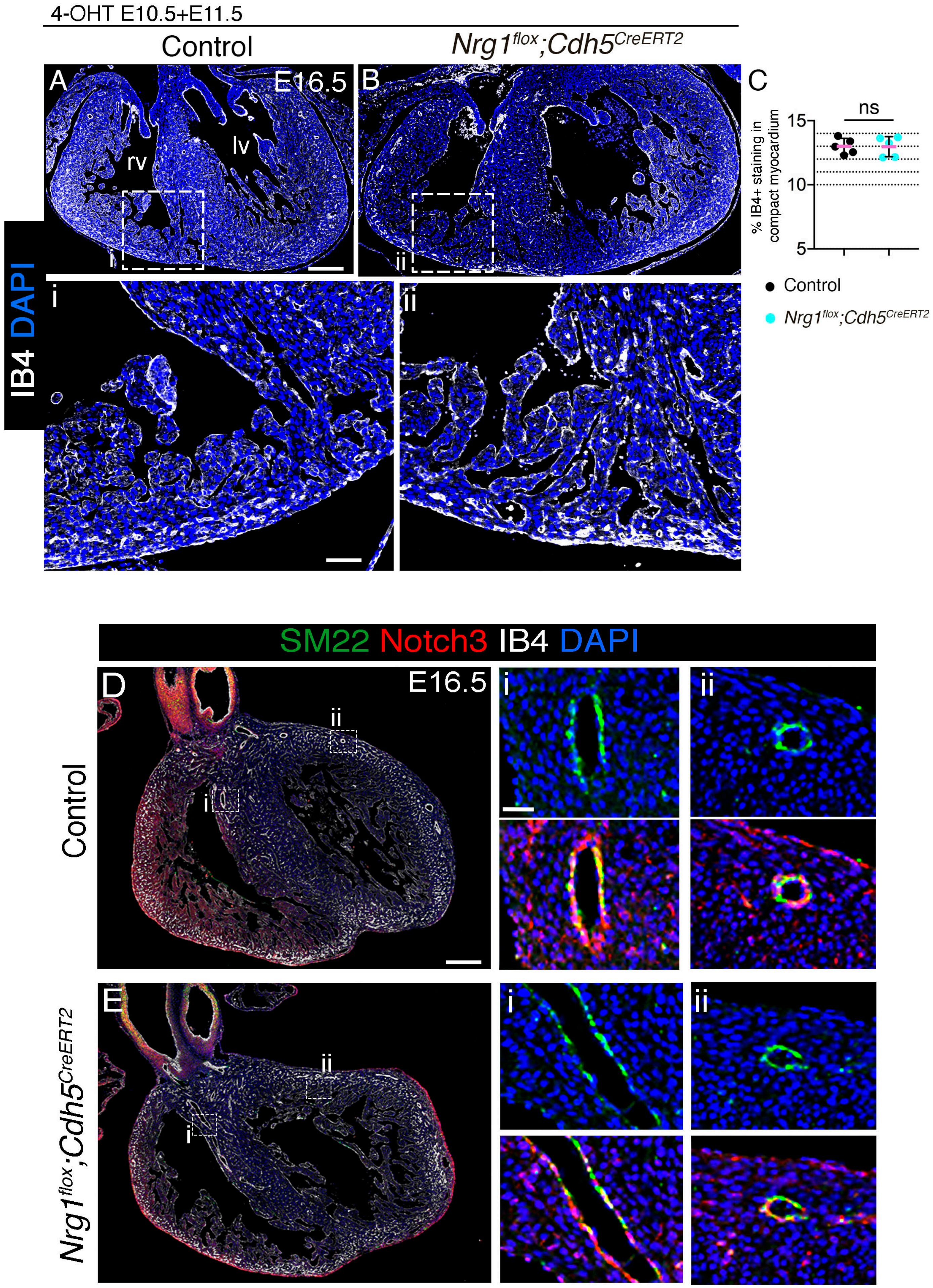


Figure S10_Grego-Bessa

**Figure S10. Coronary artery differentiation does not depend on Nrg1 signalling. (A,B)** Isolectin B4 (IB4) immunofluorescence in E16.5 control and *Nrg1$^{flox}$;Cdh5$^{CreERT}$* heart sections 4-OHT-induced at E10.5 and E11.5). DAPI counterstaining in blue**.** (i,ii) Insets showing details of right ventricle (rv). (**C**) Quantification of % of IB4+ staining in compact myocardium (n=5). ns. non-significant. Student t-test. **(D,E)** Immunofluorescence staining for SM22 (green), Notch3 (red), and IsoB4 (white) on heart sections from E16.5 control and *Nrg1$^{flox}$;Cdh5$^{CreERT}$* embryos (4-OHT induced at E10.5 and E11.5). (i,ii) Insets showing details of SM22 immunostaining and SM22/Notch2 co-immunostaining. Scale bars, 200 μm in A,B,D,E; 40 and 30 μm in insets.

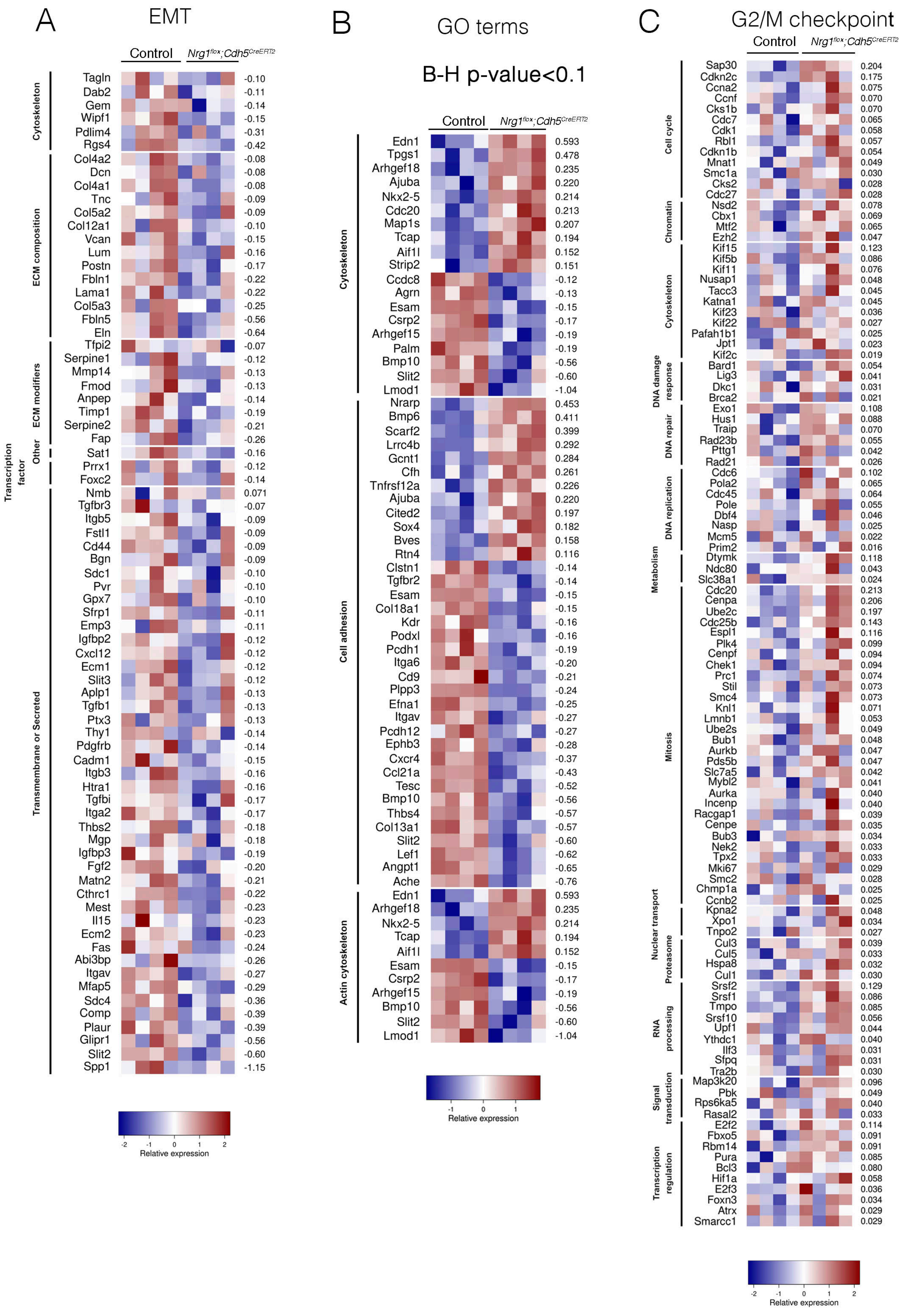


Figure S11_Grego-Bessa et al.

**Figure S11. GSEA and PANTHER analyses of E15.5 Control vs. *Nrg1$^{flox}$;Cdh5$^{CreERT2}$*. (A)** Heatmap represent relative expression values in each sample for the "leading edge" genes associated with EMT, extracted from results described in Figure 4. Genes have been manually classified into functional categories according to their description in GeneCards (https://www.genecards.org/). Numbers on the right represent logFC values. **(B)** Expression profile for differentially expressed genes annotated with GO terms "cytoskeleton organization" (GO:0007010), "cell adhesion" (GO:0007155) and "actin cytoskeleton organization" (GO:0030036). Differentially expressed genes were selected by applying a B-H adjusted p-value < 0.1 threshold. **(C)** Heatmap representing relative expression values, in each sample, for the "leading edge" genes associated with G2/M CHECKPOINT, extracted from results described in Figure 3. Genes have been manually classified into functional categories according to their description in GeneCards (https://www.genecards.org/). Numbers on the right represent logFC values.

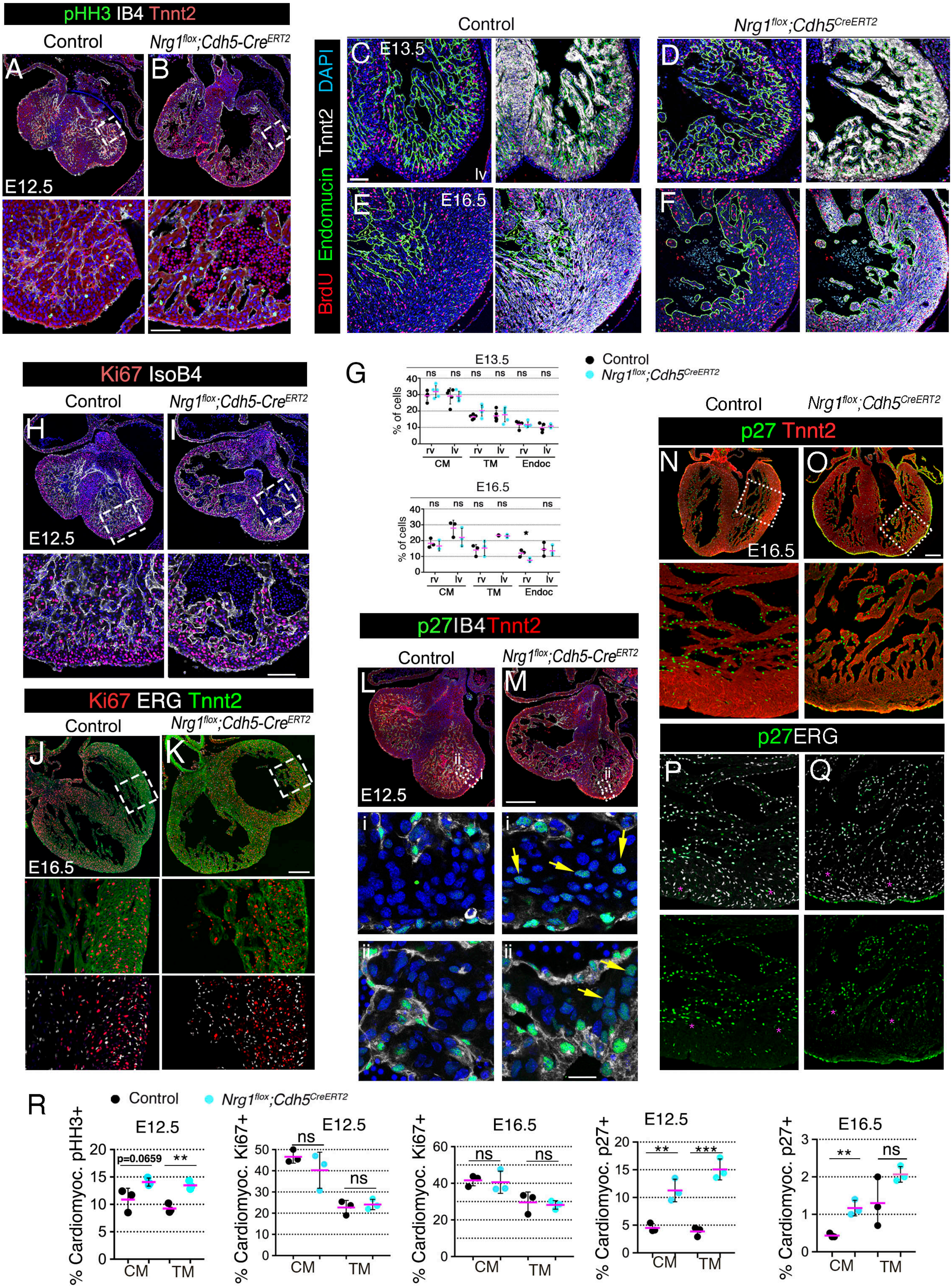


Figure S12_Grego-Bessa

**Figure S12. *Nrg1* endothelial deletion causes G2/M phase cell cycle arrest. (A,B)** Immunofluorescence staining for phospho-histone H3 (pHH3) on E12.5 control (**A**) and $Nrg1^{flox};Cdh5^{CrERT2}$ heart sections (**B**); the myocardium was immunostained for Troponin T2 (Tnnt2), the endothelium/endocardium for Isolectin B4 (IB4), and nuclei were stained with DAPI (blue). **(C,D)** BrdU immunodetection (red) on sections of control **(C,E)** and $Nrg1^{flox};Cdh5^{CrERT2}$ hearts **(D,F)** at E13.5 and E16.5. Sections were also immunostained for Endomucin (green) to mark endocardium and Tnnt2 (white) to mark myocardium; nuclei were counterstained with DAPI (blue). **(G)** Quantifications of % BrdU-positive nuclei in control and $Nrg1^{flox};Cdh5^{CrERT2}$ hearts ($n \geq 3$ per genotype) at E13.5 and E16.5. Endoc, endocardium; Student t-test. ns. non-significant. **(H-K)** Ki67 immunodetection (red) on control and $Nrg1^{flox};Cdh5^{CrERT2}$ heart sections at E12.5 **(H,I)** and E16.5 **(J,K)**. **(L-Q)** p27 immunofluorescence on control and $Nrg1^{flox};Cdh5^{CrERT2}$ heart sections at E12.5 **(L,M)** and E16.5 **(N-Q)**. The endothelium/endocardium in (**H,I,L,M**) was labeled by Isolectin B4 (IB4), and with ERG antibody in (**J,K,P,Q**); the myocardium in (**J,K,L,M,N,O**) was labeled for Tnnt2 (green or red). Sections were counterstained with DAPI (blue). **(R)** Quantification of % of cardiomyocytes pHH3-positive nuclei at E12.5, % Ki67-positive nuclei at E12.5 and E16.5, and % p27-positive nuclei at E12.5 and E16.5 in control and $Nrg1^{flox};Cdh5^{CrERT2}$ hearts (n=3 per genotype). ** $P< 0.01$; *** $P< 0.001$; ns. non-significant Student t-test. CRE activity was 4-OHT-induced at E10.5 and E11.5. rv, right ventricle; lv, left ventricle; CM, compact myocardium; TM, trabecular myocardium. Scale bars, 50 μm in A-F,H,I, L,M; 200 μm in J,K,N,O; 10 μm in insets.

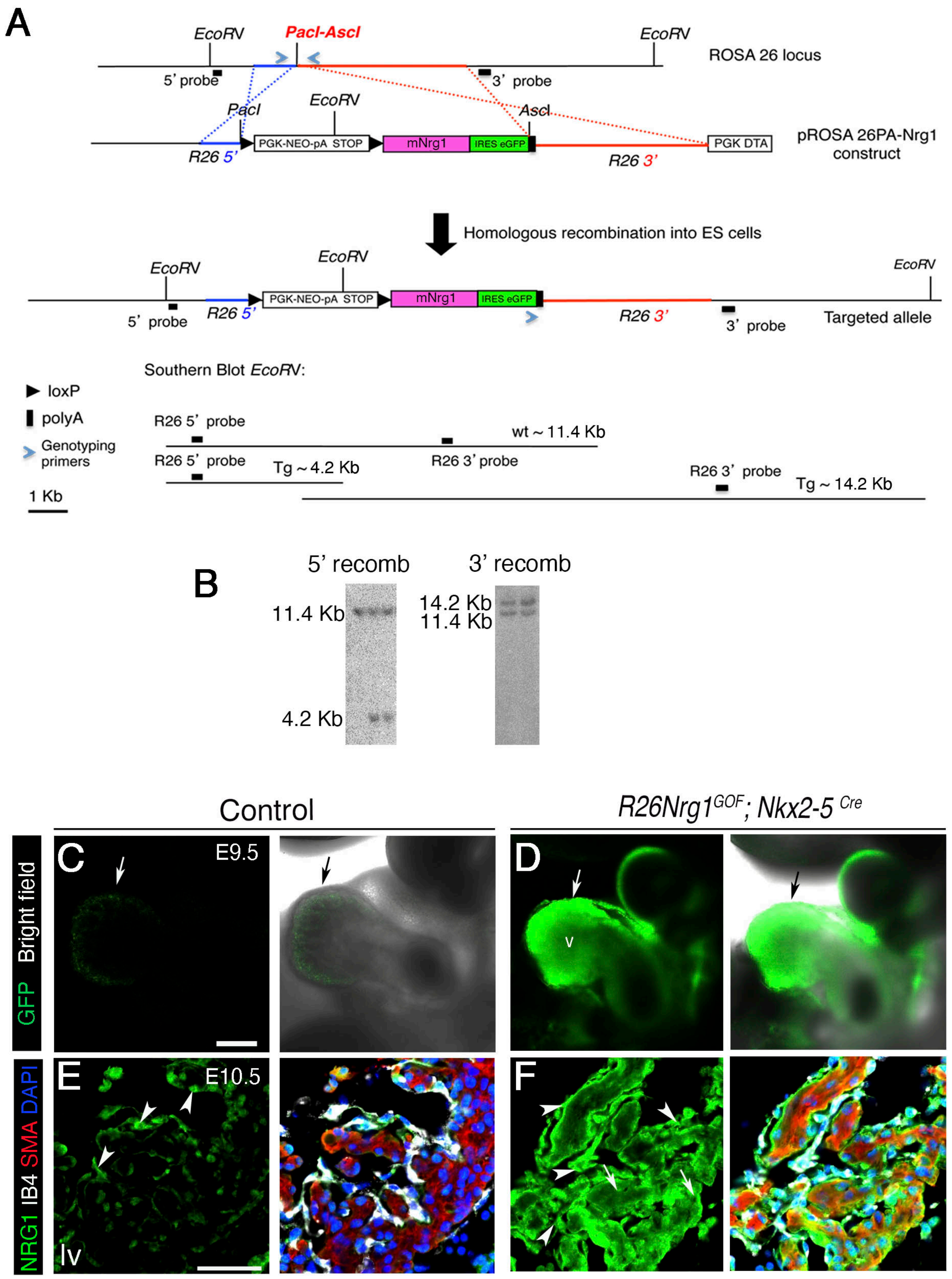


Figure S13_Grego-Bessa

**Figure S13. Generation and gene expression analysis of *R26Nrg1$^{GOF}$* mice. (A)** Gene targeting strategy in mESCs. The *floxed PGK-NEO-pA* STOP cassette followed by the *Nrg1* cDNA and an *IRES-eGFP* sequence were targeted into the *ROSA26* locus. The blue arrows indicate the position of the PCR primers used for genotyping. Black arrows indicate flanking *loxP* sites. **(B)** Positive mESC clones analyzed by Southern blot of *EcoR*V-digested genomic DNA using 5' and 3' probes that identify 4.2kb and 14.2kb transgenic bands, respectively. **(C, D)** Control and *R26Nrg1$^{GOF}$;Nkx2-5$^{Cre}$* transgenic littermate whole-mount embryos at E9.5. The left panel in each panel pair shows the transgenic GFP fluorescence signal shown separately, and the right panel in each pair shows the GFP signal superimposed on the bright-field image. The arrows mark the ventricular myocardium. **(E, F)** Nrg1 Immunofluorescence (green) in cryosections of E10.5 control and *R26Nrg1$^{GOF}$;Nkx2-5$^{Cre}$* heart sections. The arrowheads in the control panel point to normal endocardial Nrg1 expression; the arrowheads in the *R26Nrg1$^{GOF}$;Nkx2-5$^{Cre}$* panel point to ectopic myocardial Nrg1 expression. Myocardium is immunostained for smooth muscle actin (SMA; red) and endocardium (IsoB4, white); nuclei are stained with DAPI (blue). lv, left ventricle. Scale bars, 100 μm.

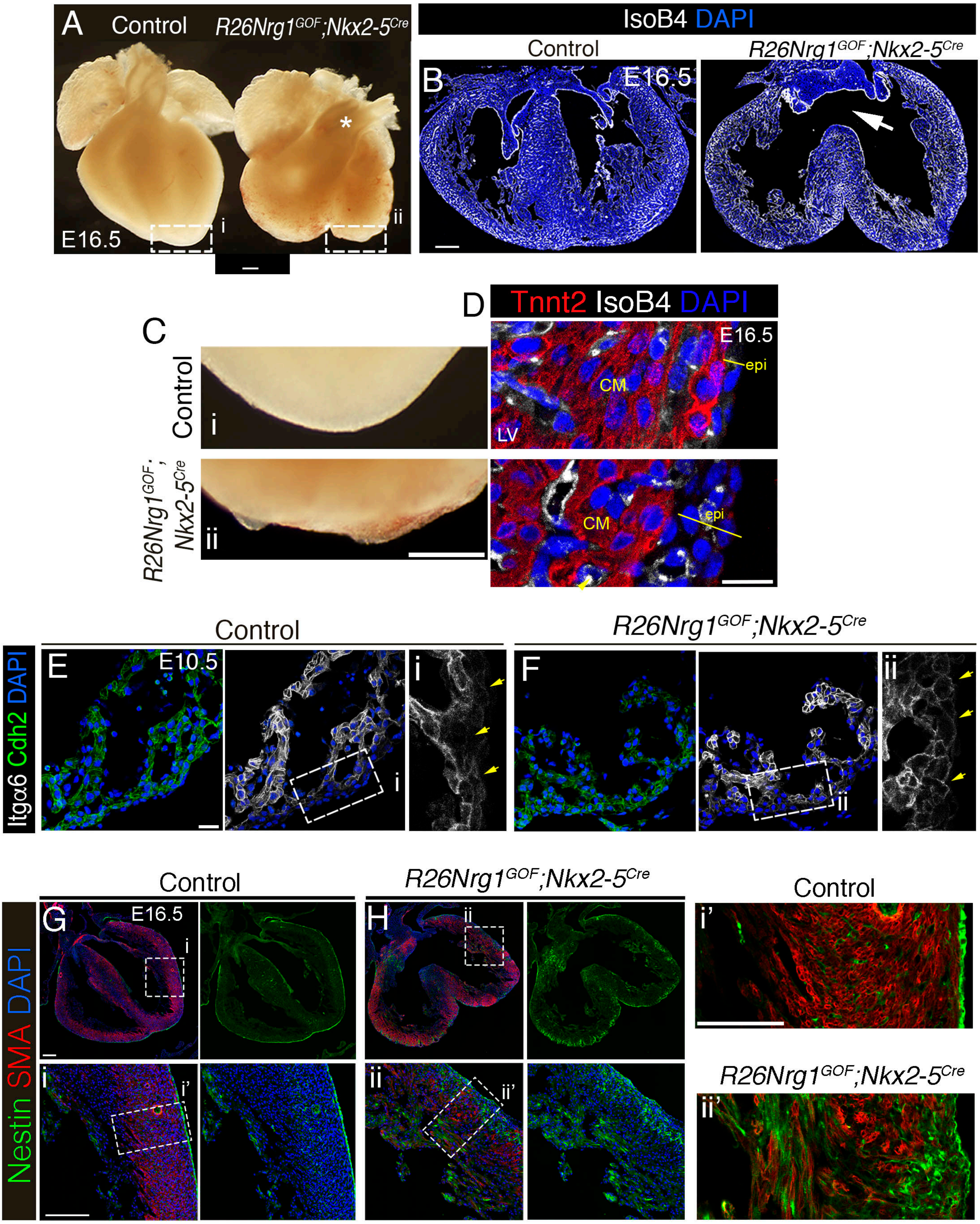


Figure S14_Grego-Bessa

**Figure S14. Ectopic Nrg1 expression disrupts heart development. (A)** Bright-field images of the ventral side of E16.5 control and *R26Nrg1*$^{GOF}$*;Nkx2-5*$^{Cre}$ hearts. **(B)** IsoB4 immunodetection in E16.5 control and *R26Nrg1*$^{GOF}$*;Nkx2-5*$^{Cre}$ heart sections (4OHT-induced at E10.5 and E11.5). DAPI counterstaining in blue**.** Arrow points to ventricular septal defect. **(C)** High-magnification views (i and ii) of the heart apexes shown in **(A)**. (**D**) High magnification of detail of left ventricle in E16.5 control and *R26Nrg1*$^{GOF}$*;Nkx2-5*$^{Cre}$ sections immunostained with Troponin T2 (Tnnt2, red), IsoB4 and DAPI counterstain. The yellow lines indicate the thickness of the epicardium (epi) as a single-cell layer in the control heart, and multilayered in the *R26Nrg1*$^{GOF}$*;Nkx2-5C*$^{re}$ heart. **(E,F)** Immunofluorescence staining of N-cadherin (Cdh2, green) and integrin α6 (Itgα6, white) on E9.5 control and *R26Nrg1*$^{GOF}$*;Nkx2-5*$^{Cre}$ heart sections. Nuclei are stained with DAPI (blue). The yellow arrows mark the outer myocardium and ectopic Itgα6 expression in compact myocardium of *R26Nrg1*$^{GOF}$*;Nkx2-5*$^{Cre}$ hearts. **(G,H)** Immunofluorescence staining for Nestin (green) on E16.5 control and *R26Nrg1*$^{GOF}$*;Nkx2-5*$^{Cre}$ heart sections; myocardium was immunostained for smooth muscle actin (SMA, red), and nuclei were stained with DAPI (blue). High magnification views i' and ii' show merged images, highlighting extensive Nestin expression in the *R26Nrg1*$^{GOF}$*;Nkx2-5*$^{Cre}$ ventricular wall**.** Scale bars, 200 µm in A, B, C, G, H; 200 µm in Ci, Cii; 20 µm in D,E,F; 100 µm in Fi',Gii'.

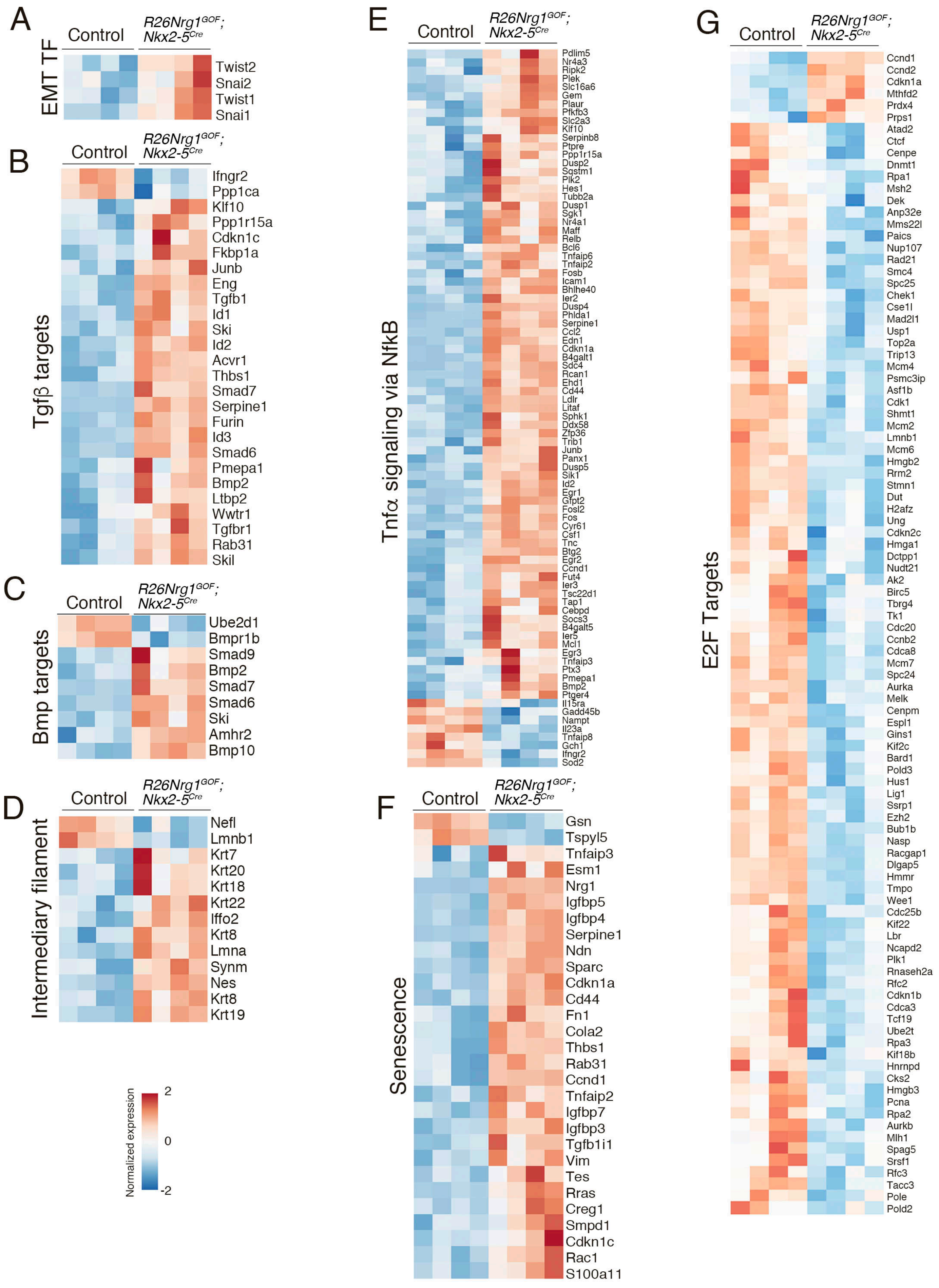


Figure S15_Grego-Bessa

**Figure S15. Heatmaps of selected DEGs and pathways in E15.5 Control vs. *Nrg1$^{flox}$;Cdh5$^{CreERT2}$*.** Heatmaps of DEGs generated in ClustVis (https://biit.cs.ut.ee/clustvis/) **(A)** Canonical EMT transcription factors. **(B)** Tgfb pathway. **(C)** Bmp pathway. **(D)** Intermediary filamets. **(E)** Tnfa-NFkB pathway. **(F)** Senescence. **(G)** E2F target genes. The color code represents normalized gene expression from -2 to +2.

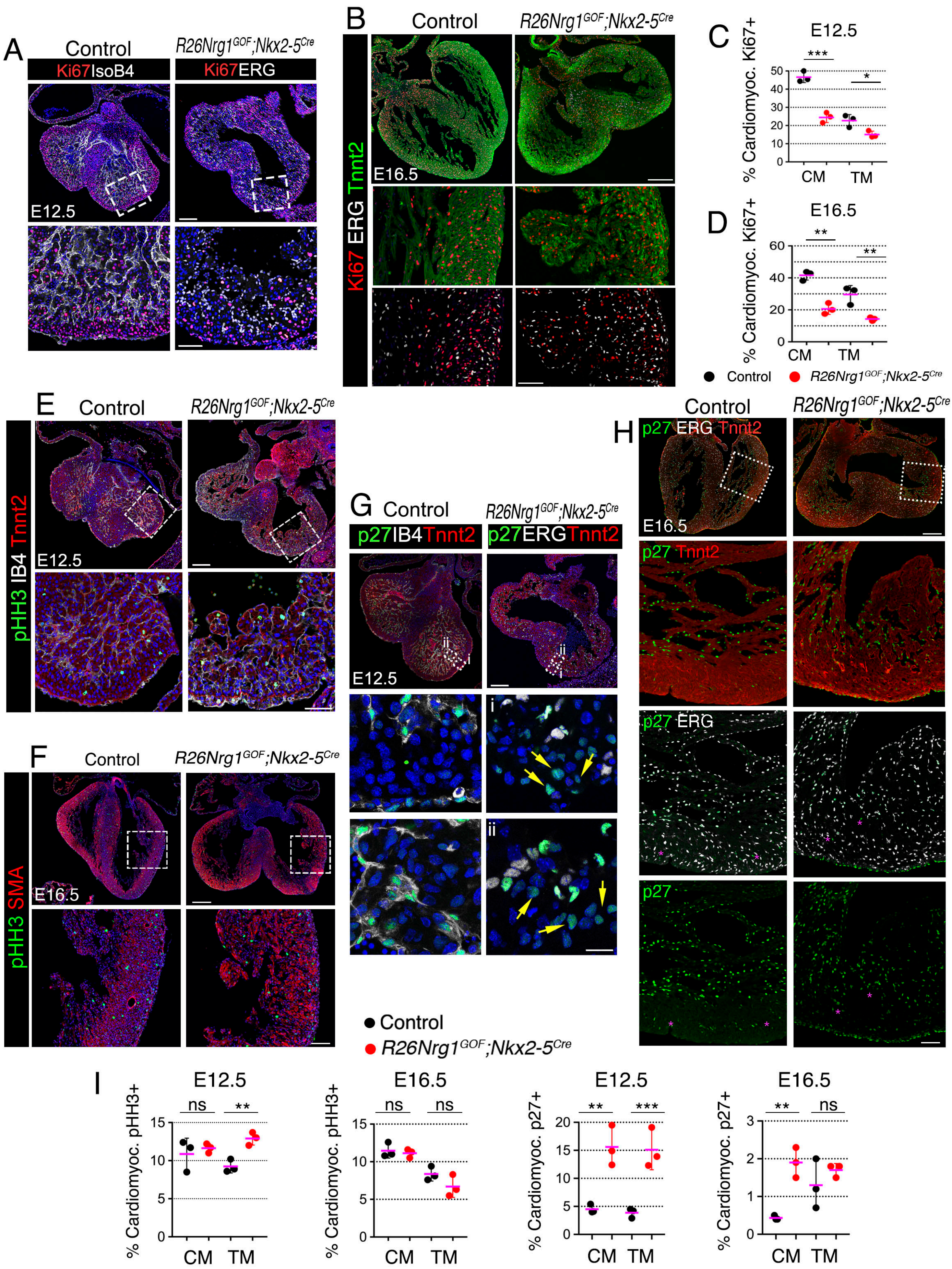


Figure S16_Grego-Bessa

**Figure S16. Ectopic cardiac Nrg1 expression causes a G1/S phase cell cycle arrest. (A,B)** Immunofluorescence staining for Ki67 antigen (red) on E12.5 control and *R26Nrg1*$^{GOF}$*;Nkx2-5*$^{Cre}$ heart sections. DAPI counterstain (blue). **(B)** Immunofluorescence staining for Ki67 antigen (red) on E16.5 control and *Nrg1*$^{flox}$*;Cdh5*$^{CrERT2}$ heart sections. Myocardium immunostained for Tnnt2 (green). Endothelium/endocardium in (**A,B**) was labeled by ERG antibody. **(C,D)** Quantification of % Ki67-positive nuclei at E12.5 (**C**) and E16.5 (**D**). **(E)** Immunofluorescence staining for phospho-histone H3 (pHH3; green) on E12.5 control and *R26Nrg1*$^{GOF}$*;Nkx2-5*$^{Cre}$ heart sections; myocardium was immunostained for Tnnt2 (red). DAPI counterstain (blue). **(F)** Immunofluorescence staining for pHH3 on E16.5 control and *R26Nrg1*$^{GOF}$*;Nkx2-5*$^{Cre}$ heart sections (green); myocardium was immunostained for SMA (red). DAPI counterstain (blue). **(G)** Immunofluorescence staining for p27 (green) on E12.5 control and *R26Nrg1*$^{GOF}$*;Nkx2-5*$^{Cre}$ heart sections. Endothelium/endocardium was labeled by Isolectin B4 (IB4) or ERG (in white). DAPI (blue) counterstain. **(H)** Immunofluorescence staining for p27 (green) on E16.5 control and *R26Nrg1*$^{GOF}$*;Nkx2-5*$^{Cre}$ heart sections. Endothelium/endocardium was labeled by ERG (white). Myocardium immunostained for Tnnt2 (red). **(I)** Quantification of % of cardiomyocytes pHH3-positive nuclei at E12.5 and E16.5, and % p27-positive nuclei at E12.5 and E16.5, in control and *R26Nrg1*$^{GOF}$*;Nkx2-5*$^{Cre}$ hearts (n=3 per genotype). * $P< 0.05$; ** $P< 0.01$; *** $P< 0.001$; ns. non-significant, Student t-test. CM, compact myocardium; TM, trabecular myocardium; Scale bars, 200 µm in B,F,H; 100 µm in A,E,G; 50 µm in insets in A,B,E,F,H; 10 µm in inset in G.

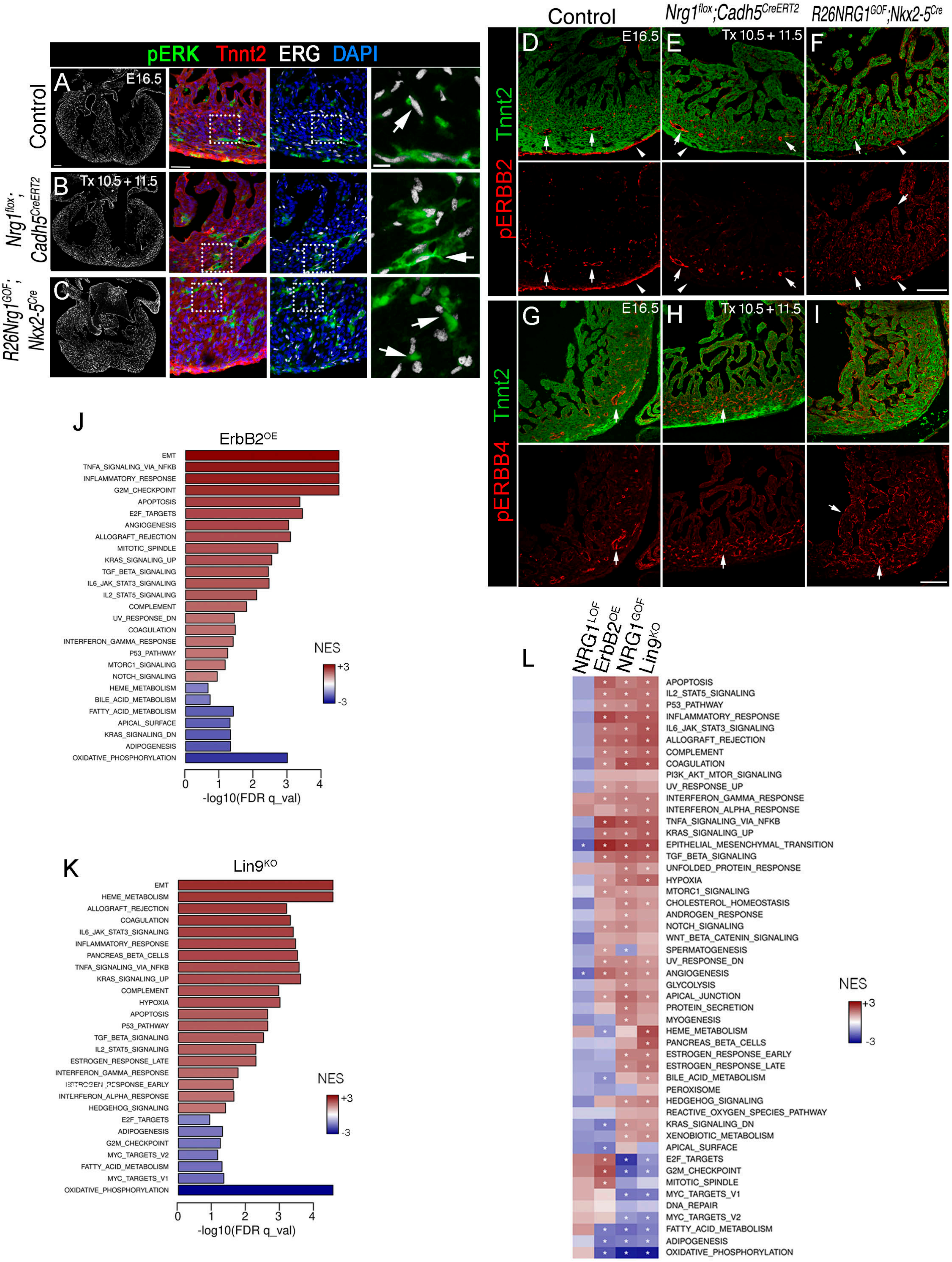


Figure S17_Grego-Bessa

**Figure S17. pErk, pErbB2 and pErbB4 expression and transcriptomic analyses in Nrg1 loss- and gain-of-function models. (A-C)** Immunofluorescence staining for pERK (green) in E16.5 control (**A**), *Nrg1$^{flox}$;Cdh5$^{CreERT2}$* (**B**), and *R26Nrg1$^{GOF}$;Nkx2-5$^{Cre}$* heart sections (**C**). Myocardium is immunostained for Tnnt2 (red), endothelial nuclei by ERG (white), and nuclei are counterstained with DAPI (blue). Arrows in the high-magnification views highlight the exclusive localization of pERK staining in endothelium. (**D-F**) Immunofluorescence staining for pErbB2 (red) on E16.5 control (**D**), *Nrg1$^{flox}$;Cdh5$^{CreERT2}$*(**E**), and *R26Nrg1$^{GOF}$;Nkx2-5$^{Cre}$* heart sections (**F**). Myocardium is immunostained for Tnnt2. *Nrg1* deletion in *Nrg1$^{flox}$;Cdh5$^{CreERT2}$* hearts was induced with 4-OHT at E10.5 and E11.5. Arrows mark pErbB2 staining in coronary endothelium; arrowheads mark staining in epicardium and sub-epicardium. **(G-I)** Immunofluorescence staining for pErbB4 (red) in E16.5 control (**G**), *Nrg1$^{flox}$;Cdh5$^{CreERT2}$* (**H**), and *R26Nrg1$^{GOF}$;Nkx2-5$^{Cre}$* heart sections (**I**). Myocardium is immunostained for TnnT2. *Nrg1* deletion in *Nrg1$^{flox}$;Cdh5$^{CreERT2}$* hearts was 4-OHT-induced at E10.5 and E11.5. Arrows mark pErbB4 staining in the coronary endothelium of all three genotypes, and additional staining in the endocardium of *R26Nrg1$^{GOF}$;Nkx2-5$^{Cre}$* hearts. Scale bars, 200 µm in A-C, insets 50µm and 10µm; 100µm in D-I. **(J)** GSEA for ERBB2$^{OE}$ vs. Control [17], against Hallmark gene sets. The barplot represents enrichment data for 27 gene sets at FDR qval < 0.25. Color scale bar indicates normalized enrichment score (NES) from -3 to +3. **(K)** GSEA for LIN9$^{KO}$ vs. Control [18], against Hallmark gene sets. The barplot represents enrichment data for 27 gene sets at FDR qval < 0.25. Color scale bar indicates NES from -3 to +3. **(L)** Comparative heatmap summarizing GSEA results for four genotypes analyzed by RNA-seq (indicated at the top of the heatmap) vs. the corresponding controls, against the Hallmark database. Blue to red colors in scale bar indicate negative to positive NES values. White asterisks indicate gene set enrichment with FDR qval<0.25.

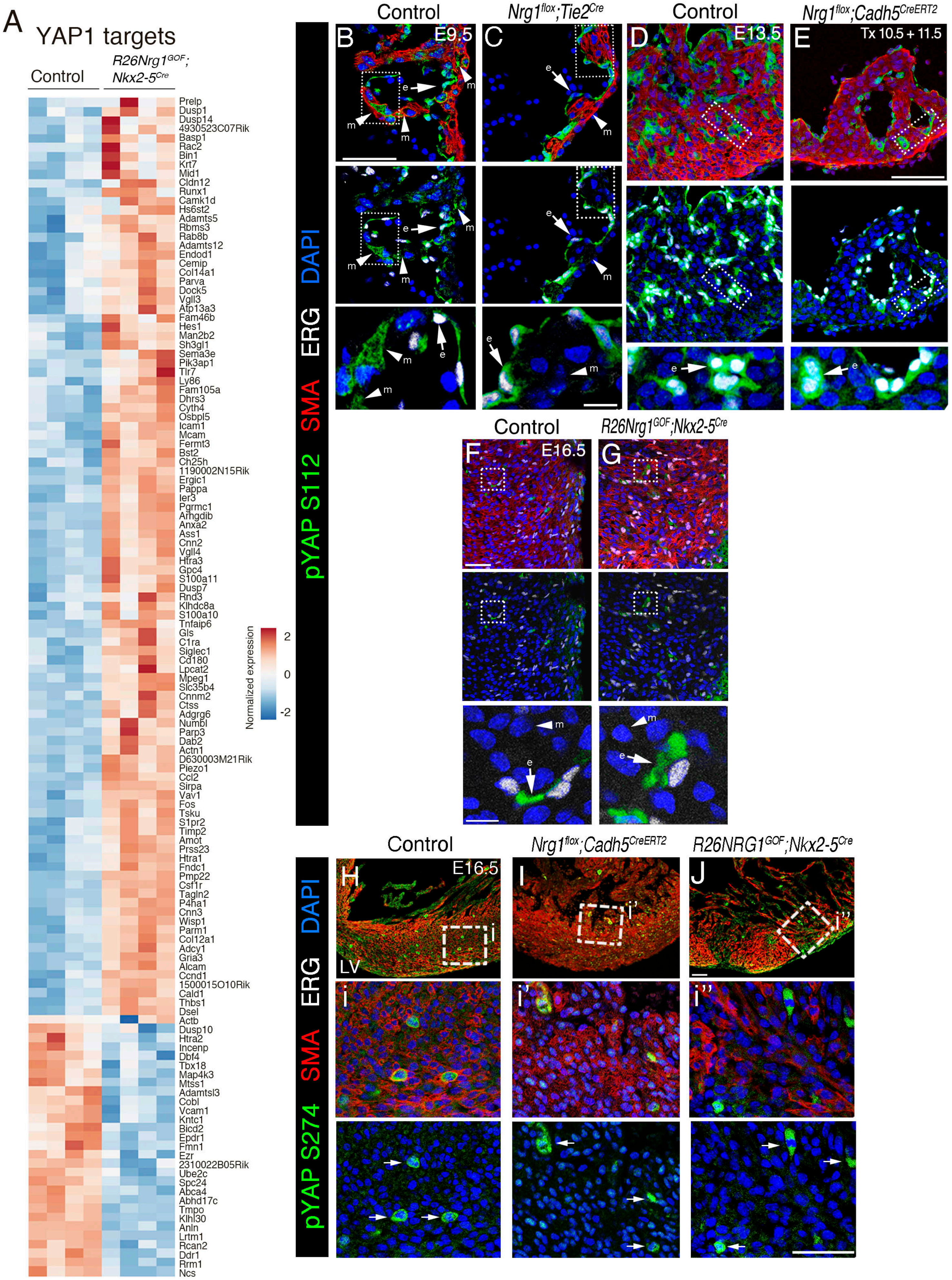


Figure S18_Grego-Bessa

**Figure S18. YAP1 target dysregulation and pYAP S112/S274 expression in Nrg1 loss-of-function and gain-of-function models.** **(A)** Heatmap of YAP1 target DEGs generated in ClustVis (https://biit.cs.ut.ee/clustvis/). The color code represents normalized gene expression from -2 to +2. **(B,C)** Detail of immunofluorescence staining for pYAP S112 (green) in E9.5 control (**B**) and *Nrg1*$^{flox}$*;Tie2*$^{Cre}$ left ventricle sections (**C**). Arrowheads mark cytoplasmic pYAP S112 staining in trabecular myocardium (m) in the control heart and the absence of myocardial pYAP S112 in the *Nrg1*$^{flox}$*;Tie2*$^{Cre}$ mutant. Arrows mark the cytoplasmic localization of pYAP S112 staining in endocardial cells (e) in both the control and the mutant. **(D,E)** Detail of immunofluorescence staining for pYAP S112 (green) on sections of E13.5 control (**D**) and *Nrg1*$^{flox}$*;Cdh5*$^{CreERT}$ hearts after 4-OHT induction at E10.5 and E11.5 (**E**). Counterstained as above. Arrows mark the exclusive localization of pYAP S112 immunostaining in endocardial cells (e) in both the control and the mutant. **(F,G)** Detail of immunofluorescence staining for pYAP S112 (green) in E16.5 control (**F**) and *R26Nrg1*$^{GOF}$*;Nkx2-5*$^{Cre}$ left ventricule sections (**G**). Arrows mark the localization of pYAP S112 immunostaining in endocardial cells in both the control and the transgenic heart; arrowheads mark the absence of staining in myocardium. In all sections the myocardium is immunostained for Tnnt2 (red) and endothelial nuclei by ERG (white), and nuclei are stained with DAPI (blue). **(H-J)** Immunofluorescence detection of pYAPS274 on E16.5 control **(H)**, *Nrg1*$^{flox}$*;Cdh5*$^{CreERT2}$ **(I)**, and *R26Nrg1*$^{GOF}$*;Nkx2-5*$^{Cre}$ **(J)** heart sections. Arrows mark pYAPS274+ mitotic nuclei. LV, left ventricle. Scale bars, 50 µm in B,D,E,F,G,H,I,J; 10 µm in high magnifications.

**Supplementary Table legends.**

**Table S1. Lethality Table**
Sheet 1: *Nrg1*$^{flox}$*;Tie2*$^{Cre}$
Sheet 2: *Nrg1*$^{flox}$*;Mesp1*$^{Cre}$
Sheet 3: *Nrg1*$^{flox}$*Cdh5*$^{CreERT2}$
Sheet 4: *R26Nrg1GOF;Nkx2-5*$^{Cre}$

**Table S2. RNAseq_E9.5_*Nrg1*$^{flox}$*;Tie2*$^{Cre}$**
Sheet 1: raw and normalized gene expression, annotations and differential expression analysis results for all genes.
Sheet 2: differentially expressed (DE) genes (adj p-val<0.05), 1219 total, 681 up-regulated (red), 538 down-regulated.
Sheet 3: GSEA Hallmark gene sets, 30 positively enriched (red), 2 negatively enriched (blue), with FDR qval<0.25.
Sheet 4: Panther enrichment results for the collection of 1219 DE genes, against the Biological Process GO term database.

**Table S3. RNAseq_E15.5_ *Nrg1*$^{flox}$*;Cdh5*$^{CreERT2}$**
Sheet 1: raw and normalized gene expression, annotations and differential expression analysis results for all genes.
Sheet 2: differentially expressed (DE) genes (adj p-val<0.05), 76 total, 44 up-regulated (red), 32 down-regulated (blue)
Sheet 3: GSEA Hallmark gene sets, 4 positively enriched (red), 10 negatively enriched (blue), with FDR qval<0.25.
Sheet 4: Panther enrichment results for the collection of 76 DE genes, against the Biological Process GO term database.

**Table S4. RNAseq_E15.5_ *R26Nrg1*$^{flox}$*;Nkx2-5*$^{Cre}$**
Sheet 1: raw and normalized gene expression, annotations and differential expression analysis results for all genes.
Sheet 2: differentially expressed (DE) genes (adj p-val<0.05), 4484 total, 2599 up-regulated (red), 1885 down-regulated (blue).
Sheet 3: GSEA Hallmark gene sets, 32 positively enriched (red), 8 negatively enriched (blue), with FDR qval<0.25.

**Table S5. PCR Primers and Antibodies**
Sheet 1: Primers
Sheet 2: Primary antibodies
Sheet 3: Secondary antibodies

**Table S6: Statistics**
Organized per Figure and Figures S

**Suppl. Movies Legends.**

**Movie S1** (from Figure 1D, E). Amira® 3D reconstruction of E10.5 control left ventricle. Z-axis is 204 µm. Whole mount (WM) IF at E10.5 (35ps) with SMA for myocardium (red) and IB4 for endocardial cells (white).

**Movie S2** (from Figure 1D, E). Amira® 3D reconstruction of E10.5 $Nrg1^{flox};Tie2^{Cre}$ left ventricle. Z-axis is 176 µm. Whole mount (WM) IF at E10.5 (35ps) with SMA for myocardium (red) and IB4 for endocardial cells (white).

**Movie S3** (from Figure 2B). Single optical sections along the Z-axis (apical to basal) of apical E8.5 control left ventricle (from 0 to 4 µm). Whole-mount phalloidin-FITC staining.

**Movie S4** (from Figure 2B). Single optical sections along the Z-axis (apical to basal) of apical E8.5 $Nrg1^{flox};Tie2^{Cre}$ left ventricle (from 0 to 4 µm). Whole-mount phalloidin-FITC staining.

**Movie S5** (from Figure 2B). Single optical sections along the Z-axis (apical to basal) of basolateral E8.5 control left ventricle (from 5 to 9 µm). Whole-mount phalloidin-FITC staining.

**Movie S6** (from Figure 2B). Single optical sections along the Z-axis (apical to basal) of basolateral E8.5 $Nrg1^{flox};Tie2^{Cre}$ left ventricle (from 5 to 9 µm). Whole-mount phalloidin-FITC staining.
